# Rapid genotyping of tilapia lake virus (TiLV) using Nanopore sequencing

**DOI:** 10.1101/2021.03.29.437503

**Authors:** Jerome Delamare-Deboutteville, Suwimon Taengphu, Han Ming Gan, Pattanapon Kayansamruaj, Partho Pratim Debnath, Andrew Barnes, Shaun Wilkinson, Minami Kawasaki, Chadag Vishnumurthy Mohan, Saengchan Senapin, Ha Thanh Dong

## Abstract

Infectious diseases represent one of the major challenges to sustainable aquaculture production. Rapid, accurate diagnosis and genotyping of emerging pathogens during early-suspected disease cases is critical to facilitate timely response to deploy adequate control measures and prevent or reduce spread. Currently, most laboratories use PCR to amplify partial pathogen genomic regions, occasionally combined with sequencing of PCR amplicon(s) using conventional Sanger sequencing services for confirmatory diagnosis. The main limitation of this approach is the lengthy turnaround time. Here, we report an innovative approach using a previously developed specific PCR assay for pathogen diagnosis combined with a new Oxford Nanopore Technologies (ONT)-based amplicon sequencing method for pathogen genotyping. Using fish clinical samples, we applied this approach for the rapid confirmation of PCR amplicon sequences identity and genotyping of tilapia lake virus (TiLV), a disease-causing virus affecting tilapia aquaculture globally. The consensus sequences obtained after polishing exhibit strikingly high identity to references derived by Illumina and Sanger methods (99.83-100%). This study suggests that ONT-based amplicon sequencing is a promising platform to deploy in regional aquatic animal health diagnostic laboratories in low and medium income countries, for fast identification and genotyping of emerging infectious pathogens from field samples within a single day.

## 1 INTRODUCTION

Aquaculture is one of the fastest growing food production sectors and is of increasing importance to global food security. This is particularly true in low income, food deficit countries, where it plays a significant role in livelihood and subsistence. However, the sustainability and expansion of the sector is hampered by disease epidemics. Endemic and emerging infectious diseases (Brummett et al., 2014), pose major animal health issues and economic losses, affecting millions of smallholders (Subasinghe et al., 2019; FAO, 2020).

Tilapia are the second most important aquaculture species (in volume) produced globally, with an industry value of $9.8 billion annually (FAO, 2020). Intensification of tilapia production has driven the emergence of diseases through the translocation of asymptomatically infected animals (Rodgers et al., 2011; Jansen et al., 2019; Dong et al., 2017a).

Rapid and accurate diagnosis of aquatic pathogens is a central pillar to any successful national aquatic animal health strategy, helping key aquaculture value chain actors to select disease-free fish broodstock, disseminate clean seeds, conduct pathogen surveillance, confirm the aetiological agent of disease outbreaks and prevent their further spread to neighbouring farms, regions and countries. This is especially important for viruses considering the lack of completely effective prophylactic treatments and vaccines for most viral pathogens of fish (Crumlish, 2017; Ninawe et al., 2017).

On suspicion of viral disease, the first recommended procedure is to demonstrate clinical pathology via simple observations of abnormal behaviours and external/internal clinical signs. Based on presumptive diagnosis using clinical signs and additional metadata collected from farmer around the disease outbreak, rapid molecular tests (such as PCR, qPCR, LAMP, or strip test kits) targeting priority pathogens can be done. Presence of viable viral particles in clinical samples can be further confirmed by culture in a permissive cell line but this can take days to weeks.

For farmed aquatic animals, molecular techniques, e.g., PCR, to confirm the presence of viral nucleic acids (DNA/RNA) are preferred because they yield much faster presumptive diagnosis. Occasionally, amplification products from semi-nested PCRs are Sanger sequenced in order to derive sequence information for genotyping, which may be used for epidemiological tracking and implementation of evidence-based biosecurity actions. Amplicon sequencing is also useful for confirmatory diagnosis to rule out possible false positive results, where less specific methods such as non-nested PCR or LAMP are used. Indeed, OIE recommends amplicon sequencing where non-nested PCR methods are employed, such as those recommended for diagnosis of Koi Herpes Virus (OIE, 2019).

Due to scarcity of sequencing facilities, with associated transport and queueing times, this process can take a few days from sample to sequence results. Unfortunately, in many low and middle-income countries (LMICs), clinical samples from disease outbreaks have to be sent overseas due to lacking of locally available sequencing capacity or limited access to specialist laboratories.

While Sanger sequencing remains the current preferred sequencing platform to produce accurate short read sequence data, it is time consuming and depends on the availability and accessibility of Sanger’s sequencing machine where needed. In addition, its analysis is somewhat laborious and may require manual inspection of the chromatogram. Second and third generation sequencing platforms such as Ion Torrent, Illumina and PacBio are extremely powerful for genomic sequencing of aquatic pathogens, but require substantial capital investment and major laboratory infrastructure. Nevertheless, they have been used to study viruses affecting global fish aquaculture, such as TiLV, piscine reovirus (PRV), piscine myocarditis virus (PMCV), salmonid alphavirus (SAV), infectious salmon anaemia virus (ISAV) (Gallagher et al., 2018; Nkili-Meyong et al., 2016).

The MinION/Flongle sequencing platform from Oxford Nanopore Technologies (ONT) offers a simple low-cost portable device for generating real-time sequence data. The low equipment cost, and particularly the lack of requirement for a well-equipped laboratory facility, makes MinION particularly attractive for genomic sequence data-driven management and control of aquatic pathogens in remote locations in LMIC. In this study, we explored the capability and advantage of ONT-based amplicon sequencing coupled with simple bioinformatics analyses for rapid and accurate consensus sequences generation for genotyping of TiLV, the causative agent of syncytial hepatitis of tilapia, a disease affecting tilapia aquaculture in over 16 countries (Taengphu et al., 2020).

TiLV is an enveloped, negative-sense, single-stranded RNA virus that contains 10 genomic segments ranging from 465 to 1641 bp, with a total genome size of 10,323 bp (Bacharach et al., 2016), encoding 14 predicted proteins (Acharya et al., 2019). The virus was recently re-classified as a new genus *Tilapinevirus*, the sole genus under the new family *Amnoonviridae* in the order *Articulavirales* (ICTV, 2020).

When new viruses (such as TiLV) emerge in aquaculture, non-validated PCR and RT-PCR methods appear very quickly after first detection of the viral diseases. Several TiLV PCR detection assays have been developed, including RT-PCR (Eyngor et al., 2014), nested RT-PCR (Kembou Tsofack et al., 2017), semi-nested RT-PCR (Taengphu et al., 2020; Dong et al., 2017b; Castañeda et al., 2020), RT-qPCR (Tattiyapong et al., 2018; Waiyamitra et al., 2018) and RT-LAMP (Phusantisampan et al., 2019; Yin et al., 2019). However with no validated OIE approved assays for TiLV, sequencing of amplicons can provide robust supporting evidence that the disease has been detected. For this study, we chose a seminested RT-PCR method (Taengphu et al., 2020) targeting TiLV segment 1, as its sensitivity is reported to be 100 times higher than a previous TiLV segment 3-based protocol (Dong et al., 2017b), and because TiLV genomic segment 1 amplicons derived from that study (Taengphu et al., 2020) have been used for genotyping of TiLV

Here, we report successful use of the semi-nested RT-PCR for diagnosis of TiLV coupled with Nanopore sequencing of amplicons for rapid identification and preliminary genotyping of TiLV. We also discuss the range of possible practical applications and implications of Nanopore sequencing, as a portable platform for robust molecular field diagnostics investigations into the origin and spread of other aquaculture pathogens of economic significance.

## 2 MATERIALS AND METHODS

### 2.1 Workflow

The diagnostic workflow from sample collection from farmed moribund fish, extraction of nucleic acid, semi-nested RT-PCR, library preparation, Nanopore sequencing and data analysis is described in Figure 1.

**FIGURE 1.**
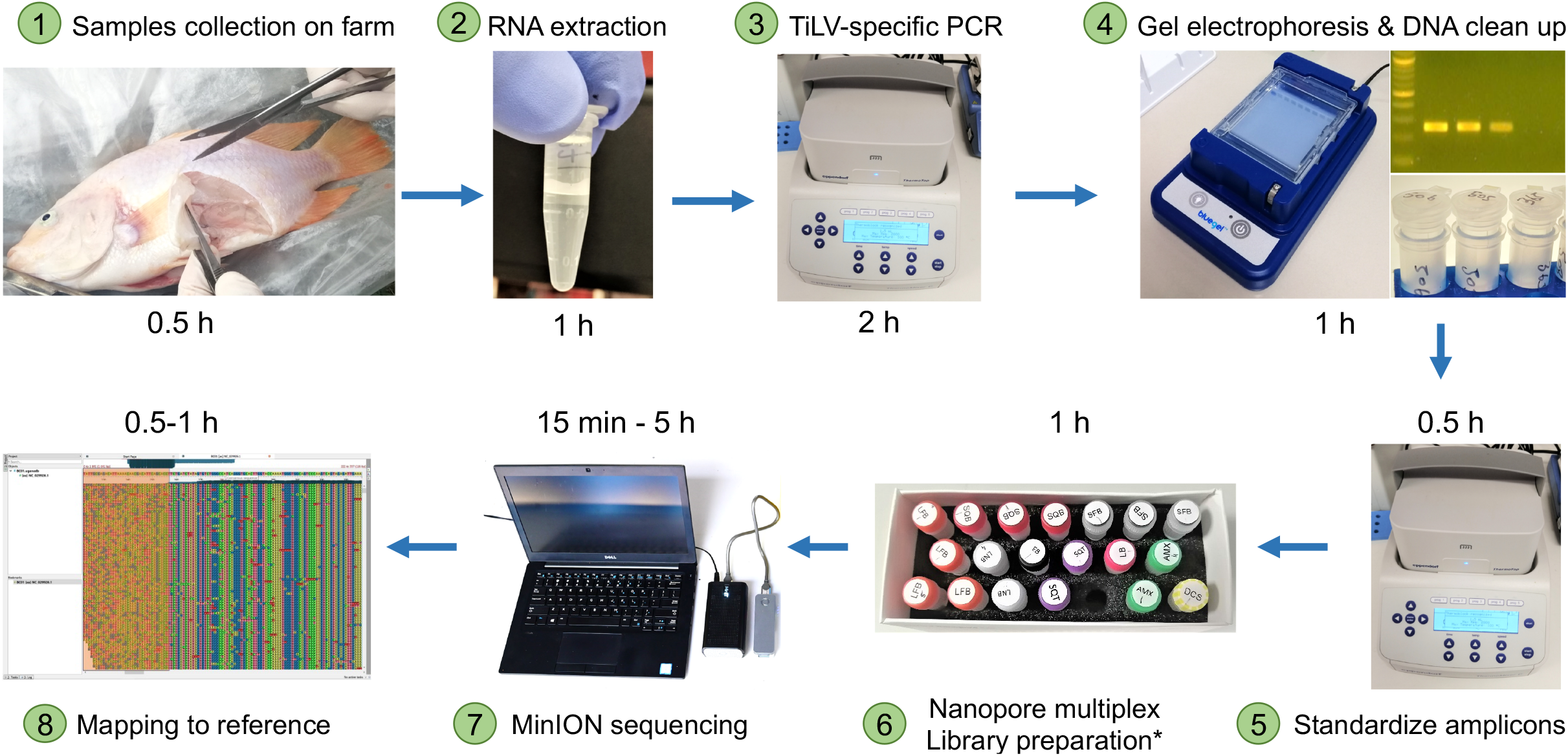
Overall workflow from sample collection of diseased fish on farm to sequence results. The entire process takes less than 12 h. * DNA repair, end-preparation, multiplex native barcode and adapter ligation

### 2.2 RNA samples and reference sequences

We used five archived RNA templates extracted from clinical Nile tilapia (*Oreochromis niloticus*, Linnaeus) and red tilapia (*Oreochromis* spp.) specimens and from E-11 permissive cell line used for TiLV propagation (Table 1). All five samples were previously confirmed to be TiLV positive. The samples were originally isolated from specimens collected in Thailand (BC01 and BC03), Bangladesh (BC02), and Peru (BC04 and BC05) as described in previous reports (Debnath et al., 2020; Taengphu et al., 2020). Table 1 also includes fourteen fulllength TiLV segment 1 reference sequences retrieved from NCBI. The NCBI reference sequences originated from tilapia specimens collected from Thailand, Bangladesh, Peru, Ecuador, Israel and the USA between 2011 and 2018, and were used for sequence alignment and phylogenetic analysis with the amplicon consensus sequences generated in this study.

**TABLE 1.**
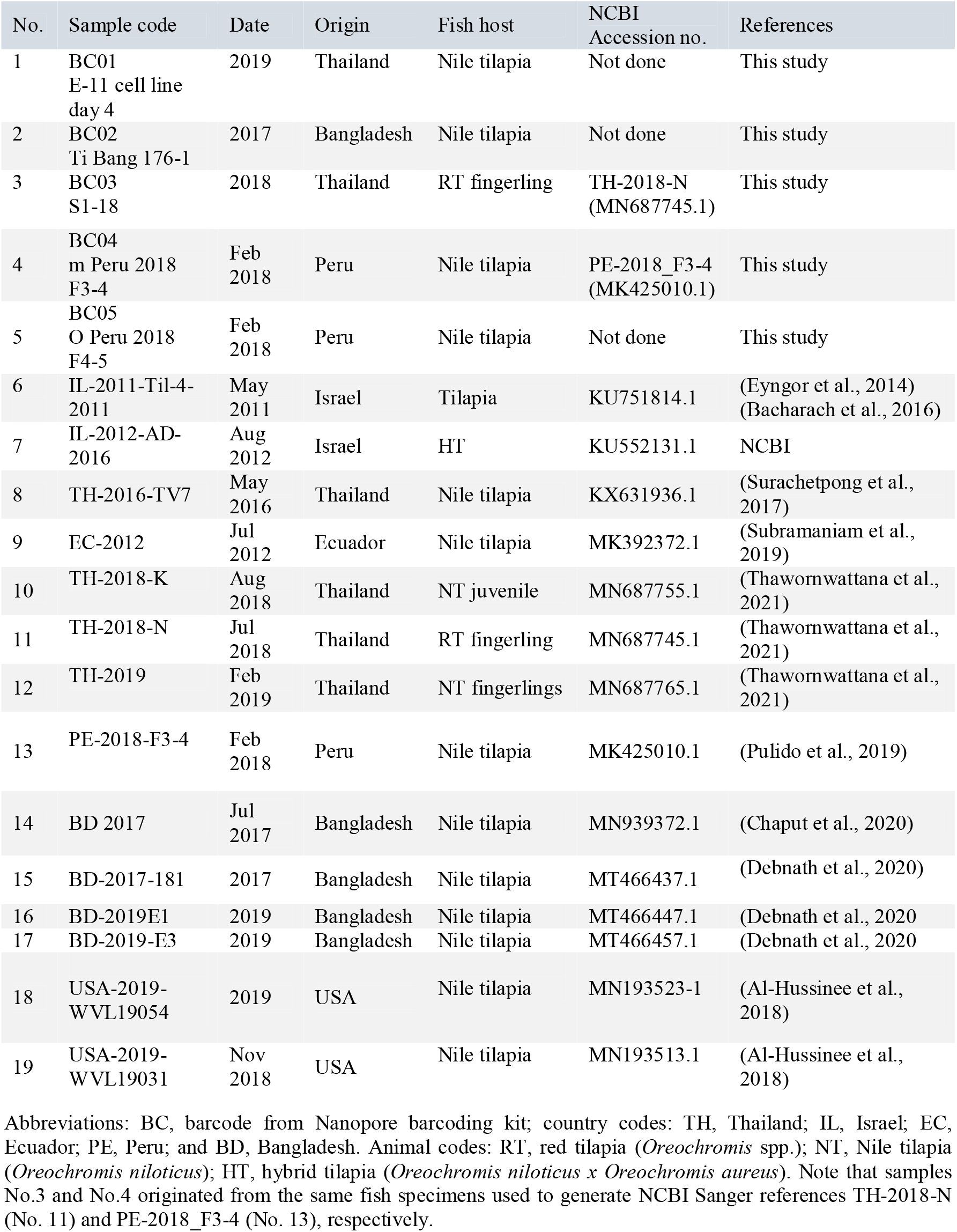
Details of TiLV samples used in this study (No. 1-5) whose genomic partial segment 1 sequences were compared with NCBI references (no. 6-19) for phylogenetic analysis

### 2.3 Semi-nested RT-PCR

Partial regions of the TiLV segment 1 genome were amplified by semi-nested RT-PCR as described previously (Taengphu et al., 2020). Five microliters of the second round PCR products were analyzed by electrophoresis on a 1% agarose gel stained with ethidium bromide solution. The remaining 20 μL reaction volume from the second round PCR was purified for each sample on a NucleoSpin Gel and PCR Clean□up column (Macherey-Nagel) and eluted with 20 μL of the kit elution buffer (5 mM Tris-HCl, pH 8.5). The purified products were quantified using Qubit dsDNA Broad Range kit (Qiagen) with a Qubit 3.0 fluorometer prior to Nanopore multiplex library preparation.

### 2.4 Library preparation of TiLV PCR products for Nanopore sequencing

To prepare the TiLV library, the ligation sequencing kit (SQK-LSK109) and the native barcoding expansion 1-12 kit (EXP-NBD104) were used according to the Oxford Nanopore Technologies (ONT) standard protocols adapted for the Flongle flow cell. We used 250 ng PCR products for each sample (BC01-BC05), one unique native barcode (BC) per sample and washed the library of pooled barcoded samples with the Short Fragment Buffer (SFB) just before the elution step at the end of the protocol. DNA concentrations were determined between each step using the Qubit assay. The prepared TiLV library was loaded as per the standard protocol onto a Flongle flow cell (FLO-FLG106)— with 29 active pores— fitted to a Flongle adapter (FLGIntSP) for MinION.

### 2.5 Data acquisition and basecalling

Control of the MinION and high accuracy base-calling data acquisition were performed offline in real-time using the stand-alone MinIT (MNT-001): a pre-configured compute module with MINKNOW software version (19.05.02). The raw Fast5 files were subsequently re-basecalled and demultiplexed using the latest Guppy version (v.4.4.1) in high accuracy mode to further improve base-calling accuracy.

### 2.6 Bioinformatics analyses for TiLV amplicons consensus sequences generation

The base-called and demultiplexed FastQ files were individually assessed using NanoStat (De Coster et al., 2018). Raw reads were aligned to a primer-trimmed TiLV Segment 1 gene region (Accession Number: MN687685.1) using Minimap2 v2.17 (-ax map-ont - secondary=no). High quality reads (qscore of 10 and above) with read length of more than 500 bp were selected for consensus generation, since they were assumed to have been generated from the sequencing of the 620 bp amplicons (first round PCR products). Briefly, the filtered reads were re-aligned to the reference sequence using Minimap2 v2.17 followed by one round of polishing with RACON v1.4.20 (-m 8 -x −6 -g −8 -w 250) and then Medaka_consensus v1.1.3 (-m r941_min_high_g360). To examine the effect of sequencing coverage (or read depth) on consensus accuracy, high-quality reads (qscore of 10 and above) ranging from 270 to 320 bp that aligned to the 274 bp amplicons (semi-nested PCR products) were randomly subsampled for 1000, 500, 100, and 50 reads and used for consensus generation as described above. Subsampling of the reads was done with seqtk v1.2 using the same initial seed number as reservoir sampling for each number of reads to be subsampled, where all reads randomly selected with equal probability.

Pair-wise nucleotide similarity of the consensus sequences against their respective reference sequences were calculated using NCBI BlastN. To obtain the number of reads sequenced over time, ‘Sequencing start time’ was extracted from every sequence identifier using grep and cut commands. The extracted data were used to generate histograms representing the number of reads generated every 5 min.

### 2.7 Alignment of TiLV segment 1 amplicon consensus sequences to public references for phylogenetic analyses

A total of 24 TiLV segment 1 sequences were used for phylogenetic analyses, including five consensus sequences derived from this study first round PCR products (620 bp), five from this study semi-nested PCR products (274 bp) and 14 full-length (1560 bp) TiLV segment 1 reference sequences retrieved from GenBank database (Table 1). The latter were trimmed to 620 bp and 274 bp. Alignments were made in Jalview (Waterhouse et al., 2009) using the web service Muscle v3.8.31 (web service) defaults parameters (Edgar, 2004). The nonaligned regions and the 5’ and 3’ primer binding sites were trimmed resulting in 577 bp and 231 bp sequences from the 620 bp and 274 bp sequences of interest, respectively. Phylogenetic trees were built in IQ-TREE (v. 1.6.12) using the Maximum Likelihood approach. The first tree using the five 577 bp consensus sequences and 14 reference sequences trimmed to 577 bp. The second tree using the five 577 bp consensus sequences trimmed to 231 bp, five original 231 bp consensus sequences and 14 reference sequences trimmed to 231 bp. Given the lack of a closely related outgroup for TILV, we opted to root the trees using the mid-point rooting method (Wohl et al., 2016) to avoid outgroup long-branches in the trees.

## 3 RESULTS

### 3.1 TiLV positive clinical samples confirmed by PCR

The segment 1 semi-nested PCR assay confirmed that the five samples used in this study (Table 1) were TiLV positive (Figure S1). Bands at 620 bp are the product of the first round RT-PCR, and bottom bands at 274 bp are the product of the second round semi-nested PCR. Samples BC01, BC02, BC03, and BC05 that produced both the 620 and 274 bp products were considered as “heavy infection” and sample BC04 that only generated a 274 bp band was considered as “light infection”. Two heavy infected samples (BC01 and BC03) yielded an additional band at around 1-1.1 kb which was derived from cross-hybridized amplified products (Figure S1) as indicated previously (Taengphu et al, 2020).

### 3.2 Sequencing output and rapid bioinformatics analyses

The sequencing run on the Flongle flow cell generated 174.69K reads with 114.99 Mb of estimated bases and 93.53 Mb base called. Depending on the sample, 517 to 964 reads were generated in the first 5 min of the run (Figure S2). Those numbers gradually decreased with reduction of available active sequencing pores to drop on average below 116 reads per sample after 4 h, 15 reads per sample after 5h and no more reads produced past 6 h of the sequencing run (Figure S2). The number of reads sequenced over time will vary depending on flow cell type (Minion vs Flongle), flow cell pore count, library preparation quality, and amplicon size. Histograms of the read length distribution—for all five samples—indicate two main peaks at 620 bp and 274 bp (Figure S3). BC01 and BC02 had a higher peak at 620 bp and BC03, BC04 and BC05 at 274 bp. Our PCR results and sequence data both confirmed the semi-quantitative nature of this (ONT)-based amplicon sequencing approach that can differentiate between heavy, medium and light TiLV infected samples (Figure S1 and S3).

Given that this is an amplicon sequencing, there is no *de novo* assembly procedure, which is typically one of the more memory-consuming step in bioinformatics. The alignment of raw Nanopore reads to the TiLV reference sequence using Minimap2 took less than 10 seconds to complete while the polishing steps consisting of RACON and MEDAKA took about 5-10 minutes per sample depending on their read depth with lower read depth leading to faster consensus generation. In this study, the entire pipeline starting from basecalled FastQ files to consensus generation, sequences alignment and phylogenetic analyses was performed on a typical office laptop (ASUS VivoBook, AMD Ryzen 5, 8 GB RAM).

### 3.3 Accurate consensus generation for TiLV identification

The average percentage identity of the adapter-trimmed and quality-filtered (qscore of 10 and above) Nanopore reads against their respective Sanger TiLV segment 1 references ranged between 92.5 and 93.2% (Table S1 and Table S2). Out of the five samples, only the Thai BC03 and Peruvian BC04 had their full length TiLV segment 1 region (1560 bp) previously Sanger sequenced: TH-2018-N and PE-2018-F3-4, respectively (Table 1 and Table 2). We confirmed 100% nucleotide identity of the 577 bp amplicon of the Thai BC03 and Peruvian BC04 consensus to their original references (Table 2A). The Thai BC01 (viral isolate from Nile tilapia infected tissue sample propagated in E-11 cell line) was also 100% identical to BC03 (isolated from red tilapia) but BC01 came from a different farm seven months later, suggesting that this variant is capable of infecting multiple species in different farming areas of Thailand.

**TABLE 2.**
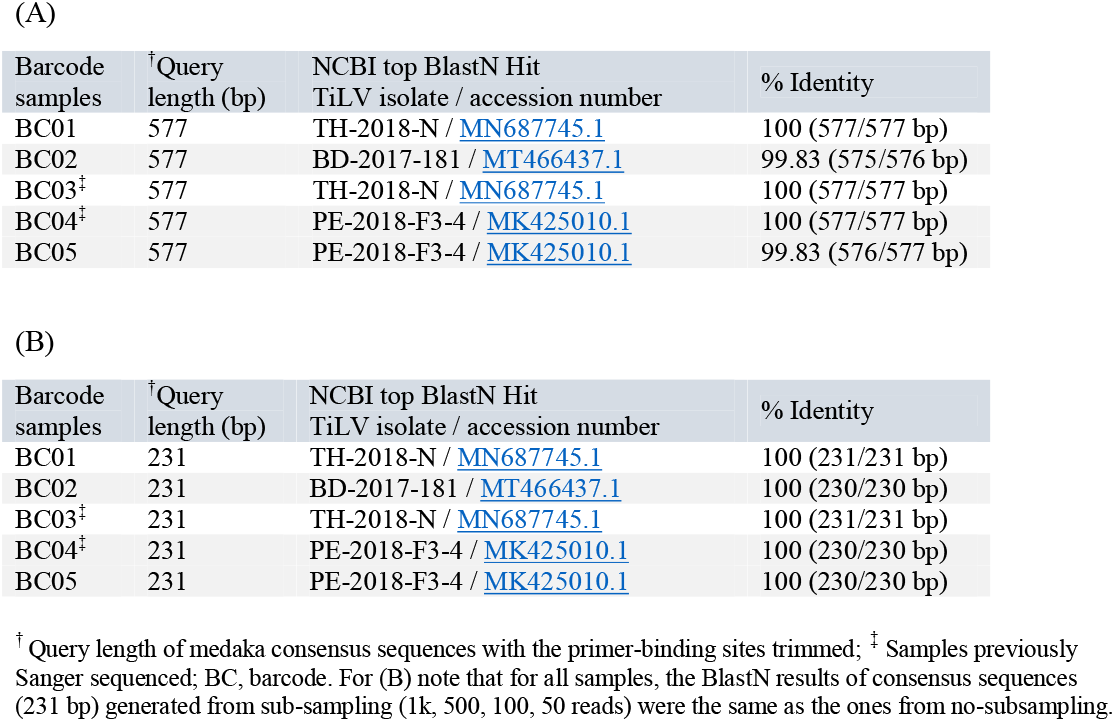
BlastN results of (A) 577 bp consensus sequences generated from the first round PCR products; (B) 231 bp consensus sequences generated from the second semi-nested round PCR products

| Barcode samples | <sup>†</sup>Query length (bp) | NCBI top BlastN Hit<br>TiLV isolate / accession number | % Identity |
| --- | --- | --- | --- |
| BC01 | 577 | TH-2018-N / MN687745.1 | 100 (577/577 bp) |
| BC02 | 577 | BD-2017-181 / MT466437.1 | 99.83 (575/576 bp) |
| BC03<sup>‡</sup> | 577 | TH-2018-N / MN687745.1 | 100 (577/577 bp) |
| BC04<sup>‡</sup> | 577 | PE-2018-F3-4 / MK425010.1 | 100 (577/577 bp) |
| BC05 | 577 | PE-2018-F3-4 / MK425010.1 | 99.83 (576/577 bp) |

The 577 bp BC02 Bangladeshi consensus sequence was 99.83% identical to BD-2017-181 (Table 2A). The single SNP (A instead of G) in position 334 (Figure 2A) was further assessed in Integrative Genomics Viewer (IGV) using BC02.medaka.bam file (read depth) with final BC02.medaka.fasta sequence. The SNP was confirmed to be amplicon-specific, partitioned between 274 and 620 amplicons (Figure 2C). Full summary of sequencing statistics for mixed amplicons (274 and 620 bp) derived from NanoStat can be found in Table S1.

**FIGURE 2.**
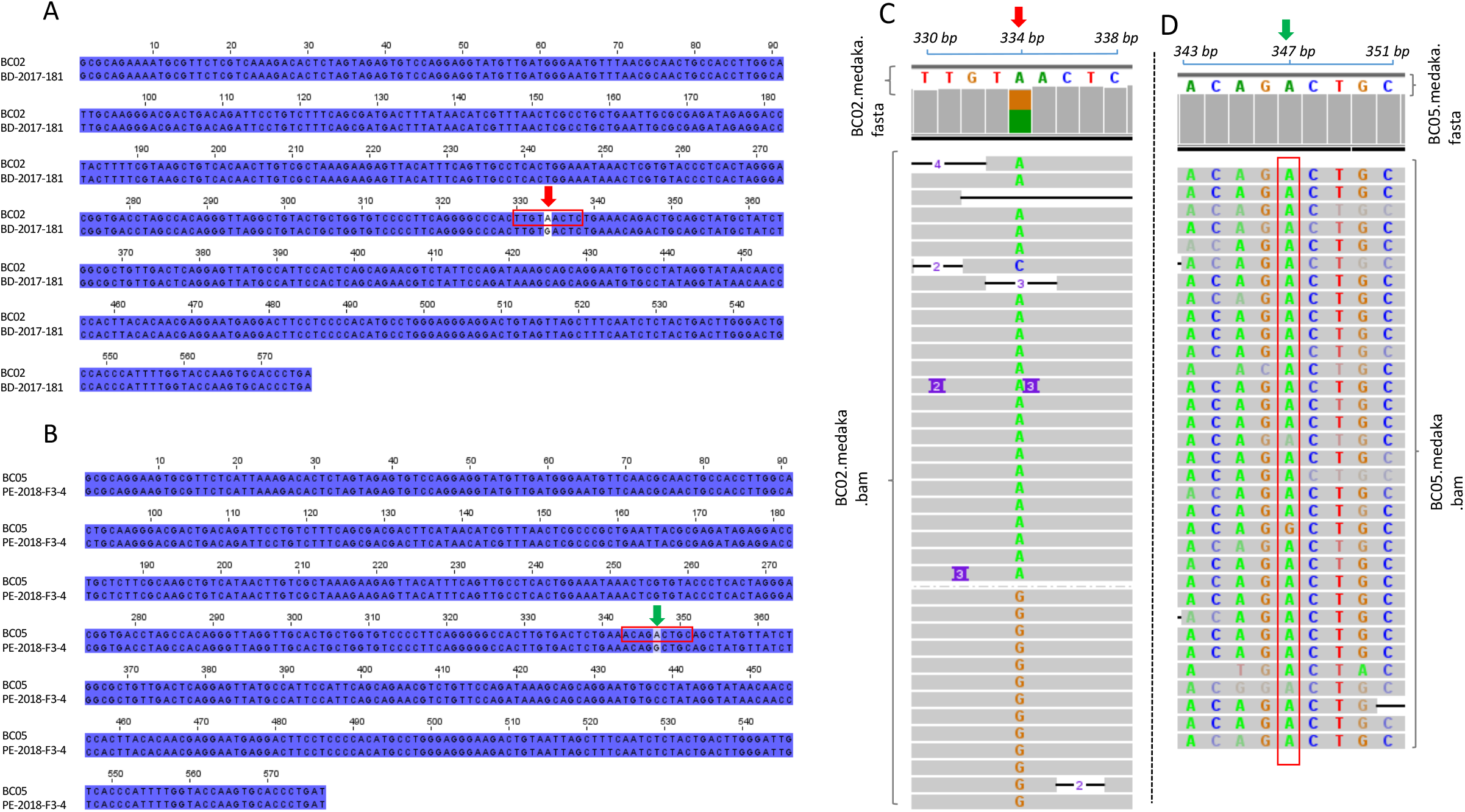
(A and B) Identification of single nucleotide polymorphisms (SNPs) using sequences alignment of TiLV segment 1 medaka consensus sequences (this study) with their closest Sanger verified references. (A) Bangladeshi BC02 consensus (576 bp) aligned with BD-2017-181 (MT466437.1) showing SNP in position 334 (red arrow); (B) Peruvian BC05 consensus (577 bp) aligned with PE-2018-F3-4 (MK425010.1) with SNP in position 347 (green arrow); (C and D) SNPs examination in Integrative Genomics Viewer (IGV) (version 2.8.10); (C) Read depth (BC02.medaka.bam file) aligned with final medaka consensus sequence (BC02.medaka.fasta file) showing the SNP is partitioned between 274 and 620 bp amplicons; (D) Read depth (BC05.medaka.bam file) aligned with final medaka consensus sequence (BC05.medaka.fasta file) confirming the SNP is real since it is identical in 97% of the reads, except for 1 homopolymer base-call error: G

A BlastN analysis of the Peruvian 577 bp BC05 consensus sequence returned 99.83% identity to PE-2018-F3-4 (Table 2A), with alignment of BC05 and PE-2018-F3-4 showing only one SNP (A instead of a G) in position 347 (Figure 2B). This SNP was confirmed in IGV, which revealed consistent base call of an Adenine (A) in the majority of the reads (BC05.consenus.bam file) with only one Guanine (G) corresponding to a homopolymer base calling error (Figure 2D). While both BC04 and BC05 were collected in 2018, they came from different farms. This indicates the presence in Peru of at least two TiLV variants at the time of sampling.

### 3.4 Sequencing coverage for reliable genotyping

Consensus sequences (231 bp) generated from the random subsampling of 1000, 500, 100 and 50 reads from the same sample are 100% similar in all cases (Table 2B). NanoStat summary statistics of sequencing output for 274 bp and sub-sampling analysis are presented in Table S2.

### 3.5 Phylogenetic analysis of TiLV segment 1 amplicon consensus

Two phylogenetic trees were generated. The first tree comparing the five 577 bp consensus sequences (this study) with NCBI reference sequences (Table 1) trimmed to 577 bp (Figure 3A). The second tree includes the same five 577 bp consensus sequences trimmed to 231 bp, with the five original 231 bp consensus sequences (this study) compared with NCBI reference sequences trimmed to 231 bp (Figure 3B).

**FIGURE 3.**
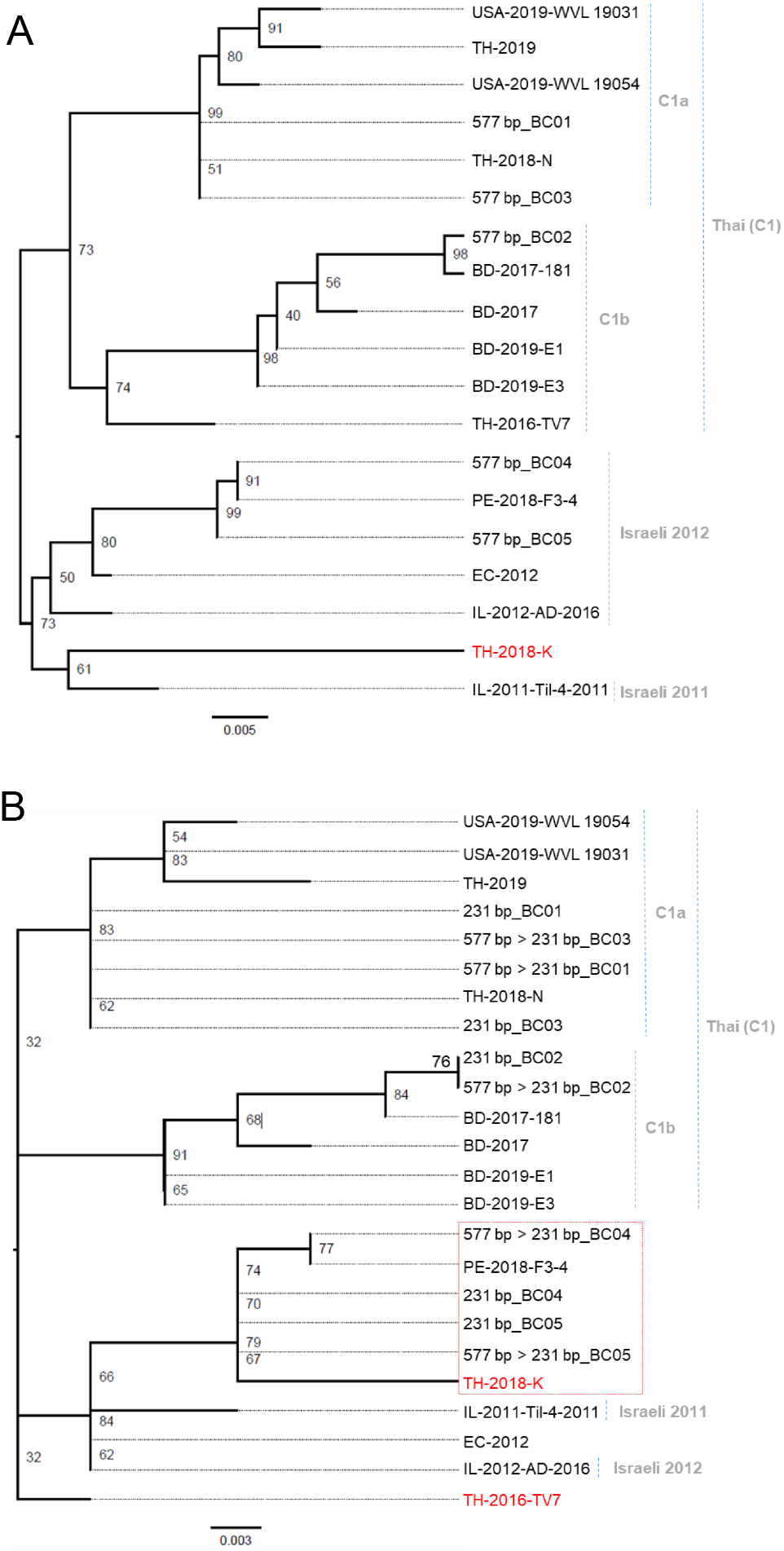
Maximum likelihood trees constructed in IQ-TREE based on the nucleotide consensus sequences alignment of short TiLV consensus (577 bp and 231 bp) with TiLV segment 1 reference sequences retrieved from GenBank database (Table 1). (A) Five 577 bp consensus sequences compared with 14 reference sequences trimmed to 577 bp. (B) five 577 bp consensus sequences trimmed to 231 bp, five original 231 bp consensus sequences compared with 14 reference sequences trimmed to 231 bp. The branch lengths indicate the number of substitutions per site, and node labels indicate bootstrap support values in percentage. Trees rooted using the mid-point rooting method

The five 577 bp consensus sequences generated in this study clustered those TiLV isolates into two separate clades, namely Thai (C1) and Israel 2012 (Figure 3A). The Thai C1 clade was divided into two sub-clades: C1a and C1b. Clade C1a contains BC01 and BC03 Thai isolates both clustering closely with TH-2018-N. Clade C1b includes BC02 that is most similar to BD-2017-181. The Israeli 2012 clade comprises BC04 and BC05 Peruvian isolates clustering with PE-2018-F3-4. IL-2011-Til-4-2011 forms a monophyletic clade outside of “Israel 2012” clade (Figure 3A). Reference sequence for TH-2018-K—when trimmed from 1560 bp to 577 bp—positions TH-2018-K outside the Thai (C1) clade (Figure 3A).

In the tree derived from alignment of 231 bp sequence data (Figure 3B), the BC01 and BC03 Thai isolates still cluster in the Thai C1a clade. The 231 bp consensus sequence of the BC02 Bangladeshi isolate still places it in the C1b Thai sub-clade but the shorter consensus sequences (231 bp) of the BC04 and BC05 Peruvian isolates, now make those two isolates more closely related to the Israeli 2011 clade (IL-2011-Til-4-2011) (Figure 3B). Trimmed reference sequences (1560 bp to 231 bp) for TH-2018-K and TH-2016-TV7 isolates now position them outside the Thai (C1) clade (Figure 3B).

## 4 DISCUSSION

The read accuracy of MinION data has been a disadvantage of the platform when compared with Sanger or Illumina sequencing. However, it has greatly improved with advances in flow cell chemistry, base-calling software and consensus accuracy. With sufficient read depth, a consensus sequence with adequate accuracy for genotyping can now be generated quickly with the right bioinformatics tools without requiring high computing capacity. With the right capacity building and training of molecular diagnosticians and aquaculture technicians, our proposed workflow and bioinformatics analytical pipeline can be adopted in targeted countries to generate similar results.

While we cannot ascertain if the SNP identified in the Bangladeshi BC02 isolate is real due to the lack of Sanger sequencing data for the same PCR product, it may be a genuine SNP variation from the viral population sequenced. BC02 was collected on the same farm and at the same time but not from the same diseased fish that was used to derive the whole genome of BD-2017-181: one of the only four publicly available TiLV segment 1-reference sequences from Tilapia in Bangladesh (Debnath et al., 2020). We know that viral RNA-dependent RNA polymerases are error-prone, with misincorporation of a wrong nucleotide estimated every 10,000-1,000,000 nucleotides polymerized depending of viral species (Sanjuán et al., 2010). This high rate of mutation comes from the lack of proofreading ability in RNA polymerases (Steinhauer et al., 1992). Given the size of the TiLV RNA genome of 10,323 bases, a mutation rate of 1 in 10,000 would mean an average of 1 mutation in every replicated genome. If a single tilapia cell is infected with TiLV and produces 10,000 new viral particles, this mutation frequency means in theory that about 10,000 new TiLV variants have been produced. This incredible high mutation rate explains why RNA viruses evolve so quickly. Viral populations even in a single infection are not homogeneous and will be mixed at any point in time during the infection. What is sequenced from the PCR is usually an amplification of the most populous variant at the time sampled with the additional stochastic effect of which templates amplify in the first few rounds of the PCR, plus the possible (but rare) misincorporation of a dNTP by the PCR polymerase early in the amplification.

Given the relatively high sequence identity (> 92.5%) at the single Nanopore read level observed for the TiLV amplicons used in this study, real-time analysis of base-called and demultiplexed Nanopore barcoded reads will allow estimation of the minimum sequencing time (or number of reads) required to achieve a positive identification, which should be occurring in just a few seconds depending on the number of samples being sequenced, flow cell pore occupancy, library preparation quality and computing capability. In this study, an amplicon read depth of 50 X is sufficient to generate a TiLV amplicon consensus sequence with high accuracy suitable for preliminary genotyping. The read depth requirement may vary depending on the sequence composition, e.g., homopolymer content that are more prone to Nanopore sequencing error. A study using Nanopore to sequence the complete genome of salmonid alphavirus (SAV1) reported similarly low sequence coverage to generate highly accurate consensus (Gallagher et al., 2018), where authors needed as little as 20 X coverage to get a consensus 99% similar to Sanger reference, while 1,000 X coverage led to 99.97% similarity.

The phylogenetic tree topology using consensus sequences of 577 bp is mostly in agreement with the literature, since it classifies the Thai and Bangladeshi consensus (BC01, BC02 and BC03) into the correct “Thai” clade. On the contrary, the Peruvian isolates (BC04 and BC05) are now more closely related to IL-2012-AD-2016 (Israeli 2012 clade)—where in other studies that used the full-length sequences (1560 bp) of TiLV segment 1—those normally cluster them into the “Israeli 2011” clade (Debnath et al., 2020; Taengphu et al., 2020). The differences observed can be explained by the different sequence lengths used between studies. Here we used shorter amplicons (231 and 577 bp) as opposed to the full-length TiLV segment 1 sequences (1560 bp) used in the two aforementioned studies, where longer sequences provide more accurate resolution.

While short amplicons seem suitable for preliminary TiLV genotyping, a recent study analyzed each individual TiLV genome segment separately, resulting in different phylogenetic trees with high estimation uncertainties (Chaput et al., 2020). The authors’ suggested exercising caution when using phylogenetic analysis to infer geographic origin and track the movement of TiLV, and recommend using whole genomes for phylogeny wherever possible. To avoid having to sequence complete viral genomes, further sequencing data may be enough to identify regions of the genome that are descriptive—similar to multi-locus sequence typing scheme used to identify prokaryote lineages. Another good example on the need for complete genomic sequences has recently been described in a study conducted by (Thawornwattana et al., 2021), which looked at eight TiLV complete genomes from Thailand collected between 2014 and 2019. Those genomes were analyzed by Bayesian inference allowing for estimation of virus evolutionary timescales, rates and global population dynamics since the early origin of TiLV. This was only possible using complete genomic sequences.

The inherent nature of segmented virus such as TiLV limits one of the main benefits of Nanopore sequencing, which is to generate a complete viral genome with a few small overlapping PCR amplified regions. Salmonid alphavirus (SAV), a ~12 kb non-segmented, single-stranded, positive-sense RNA virus is the only fish virus genome successfully sequenced by Nanopore and was confirmed for assembly accuracy against Sanger verified reference sequence (Gallagher et al., 2018). To date, the 19 complete genomes of TiLV have been sequenced by Sanger (Debnath et al., 2020; Thawornwattana et al., 2021) and, Illumina (Chaput et al., 2020; Subramaniam et al., 2019; Al-Hussinee et al., 2018), but none have been sequenced by Nanopore. To achieve this will require amplifying all 10 segments individually by RT-PCR using different primer pairs and cycling conditions and we accept that this process may be time-consuming and possibly challenging given the relatively high nucleotide divergence among TiLV strain from different lineages.

This study serves as a “proof of concept” using primers previously used to detect TiLV to reliably amplify the TiLV segment 1 gene fragment for Nanopore sequencing. That said, design of new set of universal primers to recover longer regions if not, the entire TiLV segment 1 region or more ambitiously multiple complete TiLV genome segments for Nanopore sequencing will be considered. The accuracy of Nanopore (MINION/Flongle) depends largely on the sequence composition rather than the sequence length. Generally, amplicons generated from genomic regions with longer homopolymer length will be sequenced less accurately at the single-read level. However, with sufficient read depth, a consensus with high accuracy can be generated with proper polishing step as shown in this study.

The choice of whether to sequence short amplicons, entire segment(s) or the whole genome of TiLV will depend on the specific need. For simple and rapid confirmatory PCR diagnosis results with some phylogeny inferences for preliminary genotyping, we have shown that using 274 and 620 bp amplicons from TiLV segment 1 works very well but for high-resolution epidemiological and evolutionary analyses a whole genome approach would be required.

PCR-MinION is a rapid method to generate accurate consensus sequences for TiLV identification and genotyping. This method currently takes less than 12 h from clinical sample collection to sequence results. We show that low read depth (or coverage) does not affect the accuracy of 274 bp consensus generation, hence the possibility to further reduce sequencing time. In the hands of trained and skilled end-users, this device with the specific sample preparation protocols and our analytical workflow will enable point-of-care testing and sequencing in remote locations, helping teams of governmental and supra-national institutions during disease outbreak investigations. Such application of Nanopore has been successfully applied to study human epidemics such as Ebola virus in remote areas of West Africa (Hoenen et al., 2016), the Zika virus in hard to reach regions of Brazil (Faria et al., 2016). More recently, the technology was used to sequence and identify SARS-CoV-2, the virus causing the COVID-19 pandemic (Wang et al., 2020).

In conclusion, applied to aquatic animal production systems, our approach coupled to routine diagnostic PCR, can offer a rapid and deployable mobile solution for early genotyping of TiLV and other newly emerging infectious diseases of economics importance. Genotyping provides crucial insights into the genetics of disease outbreaks and their possible origin(s). Having demonstrated that this workflow can provide genotyping information for TiLV short fragments, future work will aim at larger amplicons (> 1 kb) for finer epidemiological tracking of pathogen populations. In addition, sequencing of multiple amplicons from different samples in a single run offers scalability and the opportunity to reduce per-sample costs even further. With the deployment of portable real-time DNA sequencing platform across national reference and regional laboratories in LMIC, trained laboratory technicians will be able to genetically screen clinical samples from routine surveillance programs and disease outbreak investigations. Through genomic sequence data-driven management, competent authorities can precisely define movement controls of aquatic animals, and provide recommendations to farmers to take appropriate actions. This will minimize the introduction, and spread of TiLV and other infectious diseases of farmed aquatic animals, contributing to both economic and food security.

## Acknowledgments

This work was undertaken as part of the CGIAR Research Program on Fish Agri-Food Systems (FISH) led by WorldFish, and the CGIAR Big Data Platform Inspire Challenge 2019 led by CIAT and IFPRI. These programs are supported by contributors to the CGIAR Trust Fund. The funders provided support in the form of salary for authors [J.D.D; P.P.D; C.V.M], travels, laboratory consumables and analytical costs, but did not have any additional role in the study design, data collection and analysis, decision to publish, or preparation of the manuscript.

## Data Availability Statement

The data that support the findings of this study are available at the following links: demultiplexed FastQ files for all five samples can be found under BioProject <u>PRJNA703741</u> and BioSample accession numbers: <u>SAMN18024369</u> (BC01), <u>SAMN18024370</u> (BC02), <u>SAMN18024371</u> (BC03), <u>SAMN18024372</u> (BC04), <u>SAMN18024373</u> (BC05). The intermediate bioinformatics files (medaka.bam; medaka.bam.stats) and final consensus sequences (medaka.fasta) from partial TiLV segment 1 amplicons combined analysis (620 bp and 274 bp) and random 274 bp analysis with subsamples for 1000, 500, 100, and 50 reads, with reference fasta sequences used for both analyses can be found under Zenodo.org dataset DOI <u>10.5281/zenodo.4556414</u>.

## Author contributions

Conceptualization, J.D.D., S.S., H.T.D.; investigation, S.T., J.D.D., P.P.D., H.T.D., S.S.; formal analysis, J.D.D., H.M.G., P.K.; methodology, S.S., H.T.D., S.T., J.D.D., H.M.G.; P.K.; supervision; S.S., H.T.D., J.D.D.; writing - original draft, J.D.D., H.T.D.; review & editing, all. All authors have read and agreed to the current version of the manuscript.

## Ethics approval statement

No animal ethic approval was required since all RNA templates used in this study derived from archived samples.

## APPENDICES

### Supplementary tables legends

**TABLE S1** NanoStat summary statistics of analysis of mixed amplicons (620 and 274 bp) for each sample (BC01-05) using the full set of reads without sub-sampling. ^†^ Coverage or read depth after clustering and filtering steps; ^‡^ mean percent identity of Nanopore raw reads to each sample specific reference; ^§^ during basecalling; BC, barcode

**TABLE S2** NanoStat summary statistics of analysis of 274 bp amplicons for each sample (barcode01-05) using the full set of reads and with sub-sampling (sub1K (1000), 500, 100, or 50 reads). ^†^ Coverage or read depth after clustering and filtering steps; ^‡^ mean percent identity of Nanopore raw reads to each sample specific reference; ^§^ during basecalling; BC, barcode

### Supplementary Figures legends

**FIGURE S1** Original 1% agarose gels showing detection of partial TiLV segment 1 from five samples used in this study: (1) BC01, Thailand; (2) BC02, Bangladesh; (3) BC03, Thailand; (4) BC04, Peru; (5) BC05, Peru; other samples (a to i) were not included in this study. Gels were stained with ethidium bromide solution. M, 2-Log DNA marker (New England Biolabs); Ng, negative control. Expected band size of 620 bp and 274 bp represent amplicons from first round PCR and second round semi-nested PCR, respectively, with lanes marked +++ for heavy infection, ++ for medium infection and + for a light infection. The band marked with # on the right side of gels arose from cross hybridization of the amplified products

**FIGURE S2** Histograms of sample read counts generated every 5 min for 6 h. (A) BC01; (B) BC02; (C) BC03; (D) BC04; (E) BC05

**FIGURE S3** Distribution of sequence lengths over all sequences between 50-1399 bp from the mixed amplicons analysis (620 and 274 bp). (A) BC01; (B) BC02; (C) BC03; (D) BC04; (E) BC05. Blue arrows indicate the peaks for 274 bp amplicons and green arrows the peaks for 620 bp amplicons. Sequence length distribution obtained in FastQC High Throughput Sequence QC Report (v 0.11.9) using final medaka.bam file as inputs. Figure produced in GraphPad Prism 9.0.2; bp, base-pair

## REFERENCES

Acharya, V., Chakraborty, H. J., Rout, A. K., Balabantaray, S., Behera, B. K., & Das, B. K. (2019). Structural characterization of open reading frame-encoded functional genes from tilapia lake virus (TiLV). Molecular Biotechnology, 61, 945–957. https://doi.org/10.1007/s12033-019-00217-y

Al-Hussinee, L., Subramaniam, K., Ahasan, M. S., Keleher, B., & Waltzek, T. B. (2018). Complete genome sequence of a tilapia lake virus isolate obtained from Nile tilapia *(Oreochromis niloticus)*. Genome Announcements, 6, e00580–18. https://doi.org/10.1128/genomeA.00580-18

Bacharach, E., Mishra, N., Briese, T., Zody, M. C., Kembou Tsofack, J. E., Zamostiano, R., Berkowitz, A., Ng, J., Nitido, A., Corvelo, A., Toussaint, N. C., Abel Nielsen, S. C., Hornig, M., Del Pozo, J., Bloom, T., Ferguson, H., Eldar, A., & Lipkin, W. I. (2016). Characterization of a novel orthomyxo-like virus causing mass die-offs of tilapia. MBio, 7, e00431–16. https://doi.org/10.1128/mBio.00431-16

Brummett, R. E., Alvial, A., Kibenge, F., Forster, J., Burgos, J. M., Ibarra, R., St-Hilaire, S., Chamberlain, G. C., Lightner, D. V., Khoa, L. V., Hao, N. V., Tung, H., Loc, T. H., Reantaso, M., Wyk, P. M. V., Chamberlain, G. W., Towner, R., Villarreal, M., Akazawa, N., … Nikuli, H. L. (2014). Reducing disease risk in aquaculture. Agriculture and environmental services discussion paper; no. 9. Washington, D.C.: World Bank Group. (119 pp). Retrieved from http://documents.worldbank.org/curated/en/110681468054563438/Reducing-disease-risk-in-aquaculture

Castañeda, A. E., Feria, M. A., Toledo, O. E., Castillo, D., Cueva, M. D., & Motte, E. (2020). Detection of tilapia lake virus (TiLV) by seminested RT-PCR in farmed tilapias from two regions of Peru. Revista de Investigaciones Veterinarias del Perú (RIVEP), 31, e16158. https://doi.org/10.15381/rivep.v31i2.16158

Chaput, D. L., Bass, D., Alam, Md. M., Al Hasan, N., Stentiford, G. D., Van Aerle, R., Moore, K., Bignell, J. P., Haque, M. M., & Tyler, C. R. (2020). The segment matters: probable reassortment of tilapia lake virus (TiLV) complicates phylogenetic analysis and inference of geographical origin of new isolate from Bangladesh. Viruses, 12, 258. https://doi.org/10.3390/v12030258

Crumlish, M. (2017). Bacterial diagnosis and control in fish and shellfish. In B. Austin & A. Newaj-Fyzul (Eds.), Diagnosis and control of diseases of fish and shellfish (pp. 5–18). John Wiley & Sons, Ltd. https://doi.org/10.1002/9781119152125.ch2

De Coster, W., D’Hert, S., Schultz, D. T., Cruts, M., & Van Broeckhoven, C. (2018). NanoPack: visualizing and processing long-read sequencing data. Bioinformatics, 34, 2666–2669. https://doi.org/10.1093/bioinformatics/bty149

Debnath, P. P., Delamare Deboutteville, J., Jansen, M. D., Phiwsaiya, K., Dalia, A., Hasan, M. A., Senapin, S., Mohan, C. V., Dong, H. T., & Rodkhum, C. (2020). Two-year surveillance of tilapia lake virus (TiLV) reveals its wide circulation in tilapia farms and hatcheries from multiple districts of Bangladesh. Journal of Fish Diseases, 43, 1381–1389. https://doi.org/10.1111/jfd.13235

Dong, H. T., Ataguba, G. A., Khunrae, P., Rattanarojpong, T., & Senapin, S. (2017a). Evidence of TiLV infection in tilapia hatcheries from 2012 to 2017 reveals probable global spread of the disease. Aquaculture, 479, 579–583. https://doi.org/10.1016/j.aquaculture.2017.06.035

Dong, H. T., Siriroob, S., Meemetta, W., Santimanawong, W., Gangnonngiw, W., Pirarat, N., Khunrae, P., Rattanarojpong, T., Vanichviriyakit, R., & Senapin, S. (2017b). Emergence of tilapia lake virus in Thailand and an alternative semi-nested RT-PCR for detection. Aquaculture, 476, 111–118. https://doi.org/10.1016/j.aquaculture.2017.04.019

Edgar, R. C. (2004). MUSCLE: multiple sequence alignment with high accuracy and high throughput. Nucleic Acids Research, 32, 1792–1797. https://doi.org/10.1093/nar/gkh340

Eyngor, M., Zamostiano, R., Kembou Tsofack, J. E., Berkowitz, A., Bercovier, H., Tinman, S., Lev, M., Hurvitz, A., Galeotti, M., Bacharach, E., & Eldar, A. (2014). Identification of a novel RNA virus lethal to tilapia. Journal of Clinical Microbiology, 52, 4137–4146. https://doi.org/10.1128/JCM.00827-14

FAO. (2020). The State of World Fisheries and Aquaculture 2020: Sustainability in action. FAO, Rome. https://doi.org/10.4060/ca9229en

Faria, N. R., Sabino, E. C., Nunes, M. R. T., Alcantara, L. C. J., Loman, N. J., & Pybus, O. G. (2016). Mobile real-time surveillance of Zika virus in Brazil. Genome Medicine, 8, 97. https://doi.org/10.1186/s13073-016-0356-2

Gallagher, M. D., Matejusova, I., Nguyen, L., Ruane, N. M., Falk, K., & Macqueen, D. J. (2018). Nanopore sequencing for rapid diagnostics of salmonid RNA viruses. Scientific Reports, 8, 16307. https://doi.org/10.1038/s41598-018-34464-x

Hoenen, T., Groseth, A., Rosenke, K., Fischer, R. J., Hoenen, A., Judson, S. D., Martellaro, C., Falzarano, D., Marzi, A., Squires, R. B., Wollenberg, K. R., de Wit, E., Prescott, J., Safronetz, D., van Doremalen, N., Bushmaker, T., Feldmann, F., McNally, K., Bolay, F. K., … Feldmann, H. (2016). Nanopore sequencing as a rapidly deployable Ebola outbreak tool. Emerging Infectious Diseases, 22, 331–334. https://doi.org/10.3201/eid2202.151796

ICTV. (2020). International Committee on Taxonomy of Viruses – Virus Taxonomy: 2020 release. (2021, May 17). Retrieved from https://talk.ictvonline.org/taxonomy/

Jansen, M. D., Dong, H. T., & Mohan, C. V. (2019). Tilapia lake virus: a threat to the global tilapia industry? Reviews in Aquaculture, 11, 725–739. https://doi.org/10.1111/raq.12254

Kembou Tsofack, J. E., Zamostiano, R., Watted, S., Berkowitz, A., Rosenbluth, E., Mishra, N., Briese, T., Lipkin, W. I., Kabuusu, R. M., Ferguson, H., del Pozo, J., Eldar, A., & Bacharach, E. (2017). Detection of tilapia lake virus in clinical samples by culturing and nested reverse transcription-PCR. Journal of Clinical Microbiology, 55, 759–767. https://doi.org/10.1128/JCM.01808-16

Ninawe, A. S., Hameed, A. S. S., & Selvin, J. (2017). Advancements in diagnosis and control measures of viral pathogens in aquaculture: an Indian perspective. Aquaculture International, 25, 251–264. https://doi.org/10.1007/s10499-016-0026-9

Nkili-Meyong, A. A., Bigarré, L., Labouba, I., Vallaeys, T., Avarre, J.-C., & Berthet, N. (2016). Contribution of next-generation sequencing to aquatic and fish virology. Intervirology, 59, 285–300. https://doi.org/10.1159/000477808

OIE. (2019). Chapter 2.3.7 Infection with koi herpesvirus. In Manual of diagnostic tests for aquatic animals. OIE - World Organisation for Animal Health. Retrieved May 18, 2021, from https://www.oie.int/en/what-we-do/standards/codes-and-manuals/aquatic-manual-online-access/

Phusantisampan, T., Tattiyapong, P., Mutrakulcharoen, P., Sriariyanun, M., & Surachetpong, W. (2019). Rapid detection of tilapia lake virus using a one-step reverse transcription loop-mediated isothermal amplification assay. Aquaculture, 507, 35–39. https://doi.org/10.1016/j.aquaculture.2019.04.015

Pulido, L. L. H., Mora, C. M., Hung, A. L., Dong, H. T., & Senapin, S. (2019). Tilapia lake virus (TiLV) from Peru is genetically close to the Israeli isolates. Aquaculture, 510, 61–65. https://doi.org/10.1016/j.aquaculture.2019.04.058

Rodgers, C. J., Mohan, C. V., & Peeler, E. J. (2011). The spread of pathogens through trade in aquatic animals and their products. Revue Scientifique et Technique (International Office of Epizootics), 30, 241–256. https://doi.org/10.20506/rst.30.1.2034

Sanjuán, R., Nebot, M. R., Chirico, N., Mansky, L. M., & Belshaw, R. (2010). Viral mutation rates. Journal of Virology, 84, 9733–9748. https://doi.org/10.1128/JVI.00694-10

Steinhauer, D. A., Domingo, E., & Holland, J. J. (1992). Lack of evidence for proofreading mechanisms associated with an RNA virus polymerase. Gene, 122, 281–288. https://doi.org/10.1016/0378-1119(92)90216-C

Subasinghe, R. P., Delamare-Deboutteville, J., Mohan, C. V., & Phillips, M. J. (2019). Vulnerabilities in aquatic animal production. Revue Scientifique et Technique (International Office of Epizootics), 38, 423–436. https://doi.org/10.20506/rst.38.2.2996

Subramaniam, K., Ferguson, H. W., Kabuusu, R., & Waltzek, T. B. (2019). Genome sequence of tilapia lake virus associated with syncytial hepatitis of tilapia in an Ecuadorian aquaculture facility. Microbiology Resource Announcements, 8, e00084–19. https://doi.org/10.1128/MRA.00084-19

Surachetpong, W., Janetanakit, T., Nonthabenjawan, N., Tattiyapong, P., Sirikanchana, K., & Amonsin, A. (2017). Outbreaks of tilapia lake virus infection, Thailand, 2015–2016. Emerging Infectious Diseases, 23, 1031–1033. https://doi.org/10.3201/eid2306.161278

Taengphu, S., Sangsuriya, P., Phiwsaiya, K., Debnath, P. P., Delamare-Deboutteville, J., Mohan, C. V., Dong, H. T., & Senapin, S. (2020). Genetic diversity of tilapia lake virus genome segment 1 from 2011 to 2019 and a newly validated semi-nested RT-PCR method. Aquaculture, 526, 735423. https://doi.org/10.1016/j.aquaculture.2020.735423

Tattiyapong, P., Sirikanchana, K., & Surachetpong, W. (2018). Development and validation of a reverse transcription quantitative polymerase chain reaction for tilapia lake virus detection in clinical samples and experimentally challenged fish. Journal of Fish Diseases, 41, 255–261. https://doi.org/10.1111/jfd.12708

Thawornwattana, Y., Dong, H. T., Phiwsaiya, K., Sangsuriya, P., Senapin, S., & Aiewsakun, P. (2021). Tilapia lake virus (TiLV): genomic epidemiology and its early origin. Transboundary and Emerging Diseases, 68, 435–444. https://doi.org/10.1111/tbed.13693

Waiyamitra, P., Tattiyapong, P., Sirikanchana, K., Mongkolsuk, S., Nicholson, P., & Surachetpong, W. (2018). A TaqMan RT-qPCR assay for tilapia lake virus (TiLV) detection in tilapia. Aquaculture, 497, 184–188. https://doi.org/10.1016/j.aquaculture.2018.07.060

Wang, M., Fu, A., Hu, B., Tong, Y., Liu, R., Liu, Z., Gu, J., Xiang, B., Liu, J., Jiang, W., Shen, G., Zhao, W., Men, D., Deng, Z., Yu, L., Wei, W., Li, Y., & Liu, T. (2020). Nanopore targeted sequencing for the accurate and comprehensive detection of SARS-CoV-2 and other respiratory viruses. Small, 16, 2002169. https://doi.org/10.1002/smll.202002169

Waterhouse, A. M., Procter, J. B., Martin, D. M. A., Clamp, M., & Barton, G. J. (2009). Jalview Version 2—a multiple sequence alignment editor and analysis workbench. Bioinformatics, 25, 1189–1191. https://doi.org/10.1093/bioinformatics/btp033

Wohl, S., Schaffner, S. F., & Sabeti, P. C. (2016). Genomic analysis of viral outbreaks. Annual Review of Virology, 3, 173–195. https://doi.org/10.1146/annurev-virology-110615-035747

Yin, J., Wang, Q., Wang, Y., Li, Y., Zeng, W., Wu, J., Ren, Y., Tang, Y., Gao, C., Hu, H., & Bergmann, S. M. (2019). Development of a simple and rapid reverse transcription–loopmediated isothermal amplification (RT-LAMP) assay for sensitive detection of tilapia lake virus. Journal of Fish Diseases, 42, 817–824. https://doi.org/10.1111/jfd.12983

