## Supplementary figures and images for "Rapid genotyping of tilapia lake virus (TiLV) using Nanopore sequencing"

### Figure S1

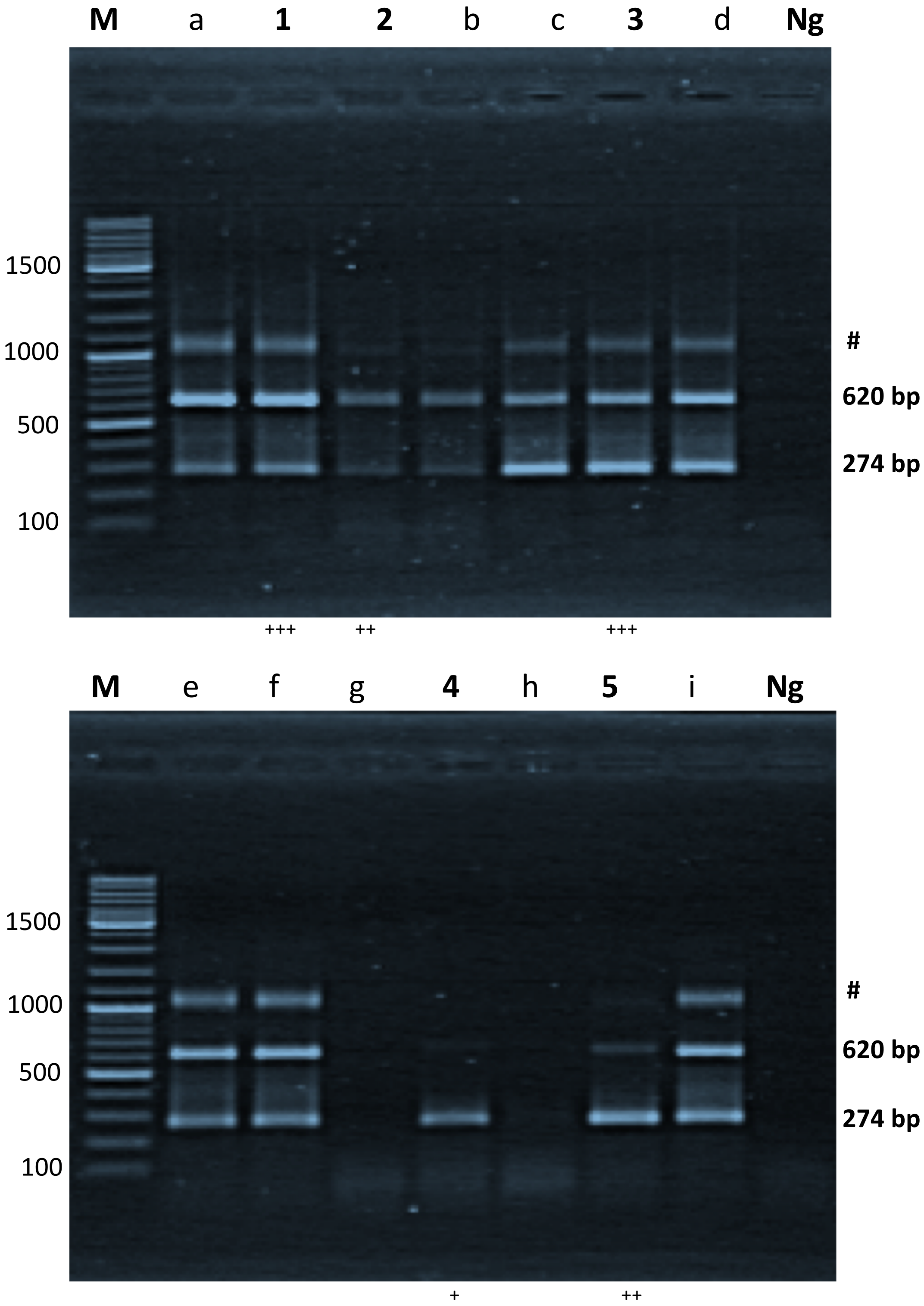

### Figure S2

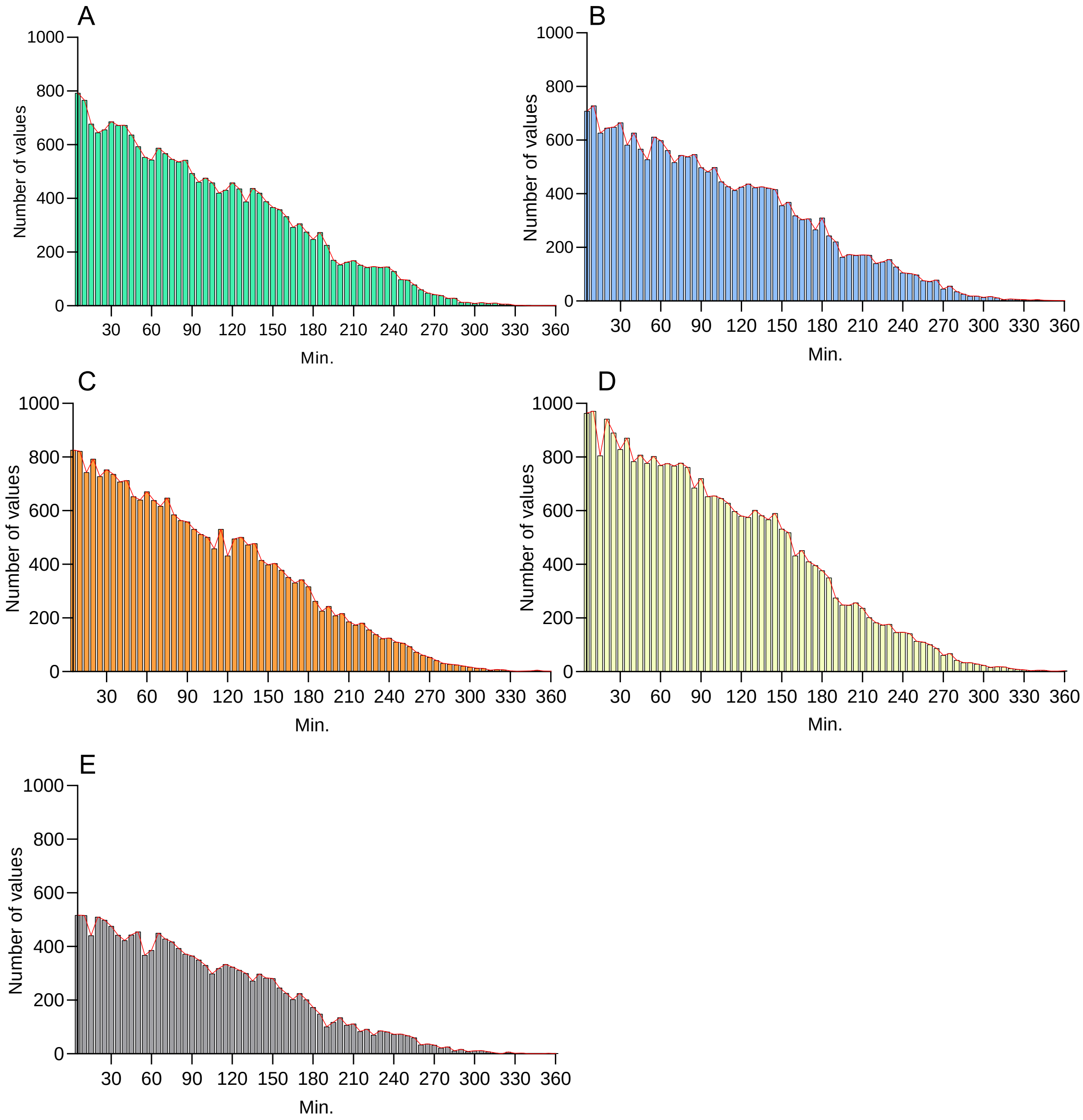

### Figure S3

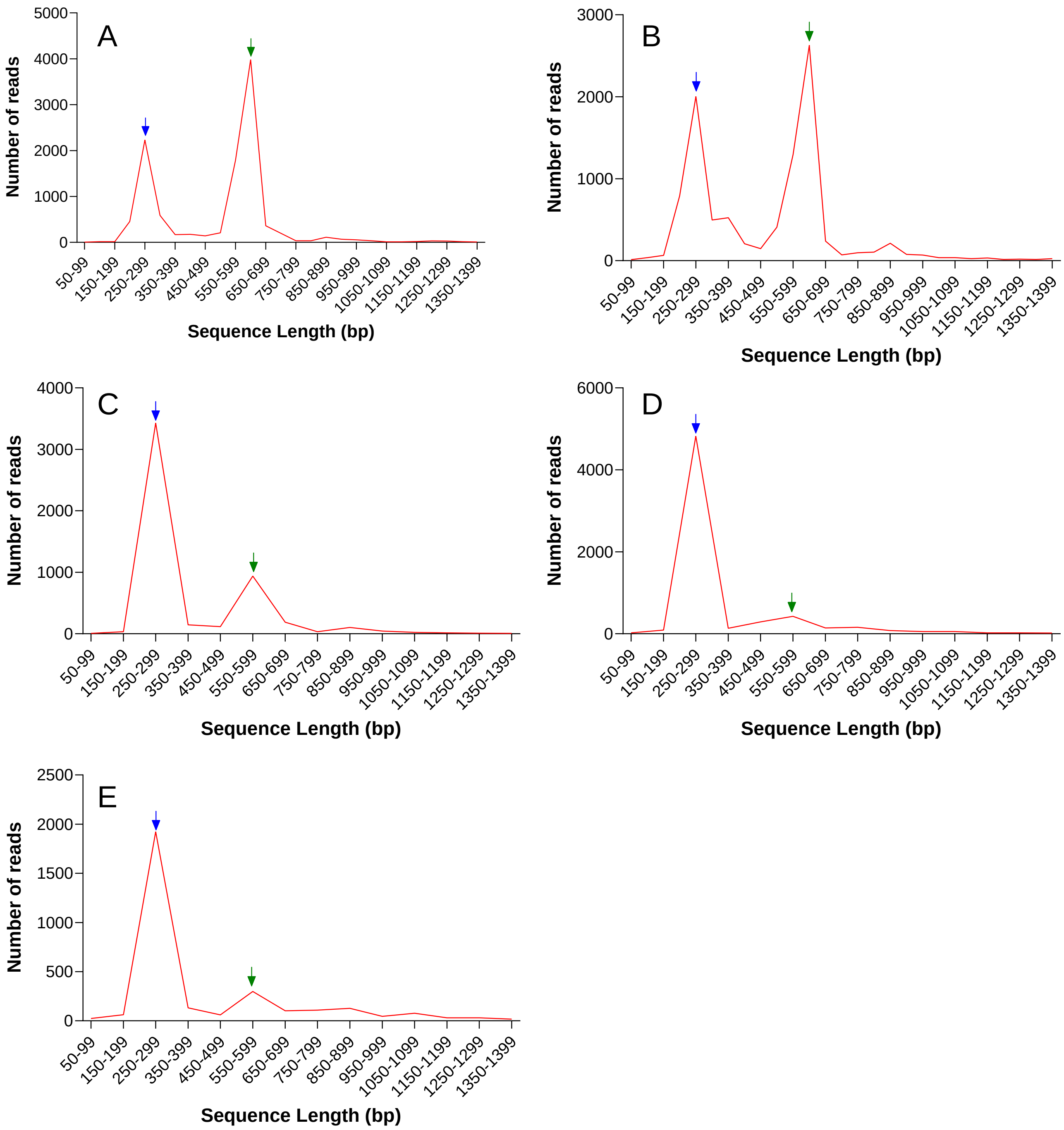
