## Supplementary material for "Rapid genotyping of tilapia lake virus (TiLV) using Nanopore sequencing": Table S1

**TABLE S1** NanoStat summary statistics of analysis of mixed amplicons (620 and 274 bp) for each sample (BC01-05) using the full set of reads without sub-sampling. ^†^ Coverage or read depth after clustering and filtering steps; ^‡^ mean percent identity of Nanopore raw reads to each sample specific reference; ^§^ during basecalling; BC, barcode

|  | **BC01** | **BC02** | **BC03** | **BC04** | **BC05** |
| --- | --- | --- | --- | --- | --- |
| number of reads (coverage) ^†^ | 10671 | 9718 | 9188 | 12307 | 7412 |
| Total bases | 5511635 | 4860840 | 4023794 | 4729660 | 3191278 |
| Total bases aligned | 4784228 | 3995675 | 3329554 | 3250112 | 2098572 |
| Fraction of bases aligned | 0.9 | 0.8 | 0.8 | 0.7 | 0.7 |
| Median read length | 596 | 577 | 323 | 277 | 278 |
| Mean read length | 516.5 | 500.2 | 437.9 | 384.3 | 430.6 |
| STDEV read length | 197.1 | 224.3 | 207.7 | 213.3 | 270.7 |
| Read length N50 | 604 | 602 | 593 | 459 | 537 |
| Mean percent identity ^‡^ | 92.6 | 92.5 | 92.9 | 93.2 | 93.1 |
| Mean percent raw reads error ^§^ | 7.4 | 7.5 | 7.1 | 6.8 | 6.9 |
| Median percent identity | 92.8 | 92.8 | 93.1 | 93.5 | 93.5 |
| Mean read quality | 12.1 | 12.1 | 12.2 | 12.3 | 12.2 |
| Median read quality | 12 | 12 | 12 | 12.1 | 12.1 |
| Top 5 longest reads and their mean basecall (quality score) |  |  |  |  |  |
| 1 | 2333 (10.1) | 2342 (10.9) | 2086 (11.9) | 2258 (10.1) | 2437 (10.1) |
| 2 | 2084 (13.0) | 2317 (11.6) | 2050 (11.3) | 2196 (11.7) | 2296 (10.1) |
| 3 | 1919 (12.2) | 2049 (12.1) | 2009 (10.1) | 2052 (10.8) | 2116 (10.1) |
| 4 | 1844 (13.6) | 2036 (10.2) | 1982 (10.5) | 2025 (11.4) | 2039 (14.0) |
| 5 | 1752 (10.9) | 2005 (10.9) | 1957 (11.1) | 1951 (10.3) | 1917 (11.6) |
| Top 5 highest mean basecall quality scores and their read lengths |  |  |  |  |  |
| 1 | 19.2 (183) | 19.4 (70) | 19.4 (251) | 20.4 (254) | 20.1 (90) |
| 2 | 18.7 (277) | 18.8 (229) | 18.8 (255) | 20.2 (273) | 20.1 (246) |
| 3 | 18.6 (246) | 18.8 (212) | 18.8 (272) | 19.9 (273) | 20.0 (249) |
| 4 | 18.3 (239) | 18.7 (237) | 18.3 (181) | 19.6 (252) | 18.8 (272) |
| 5 | 18.2 (274) | 18.7 (277) | 18.1 (274) | 19.5 (272) | 18.5 (272) |
| Number, percentage and megabases of reads above quality cutoffs |  |  |  |  |  |
| >Q5 | 10671 (100.0%) 5.5Mb | 9718 (100.0%) 4.9Mb | 9188 (100.0%) 4.0Mb | 12307 (100.0%) 4.7Mb | 7412 (100.0%) 3.2Mb |
| >Q7 | 10669 (100.0%) 5.5Mb | 9712 (99.9%) 4.9Mb | 9188 (100.0%) 4.0Mb | 12295 (99.9%) 4.7Mb | 7405 (99.9%) 3.2Mb |
| >Q10 | 10626 (99.6%) 5.5Mb | 9617 (99.0%) 4.8Mb | 9120 (99.3%) 4.0Mb | 12089 (98.2%) 4.7Mb | 7251 (97.8%) 3.2Mb |
| >Q12 | 5318 (49.8%) 2.7Mb | 4785 (49.2%) 2.2Mb | 4696 (51.1%) 1.9Mb | 6541 (53.1%) 2.3Mb | 3814 (51.5%) 1.4Mb |
| >Q15 | 299 (2.8%) 0.1Mb | 273 (2.8%) 0.1Mb | 393 (4.3%) 0.1Mb | 721 (5.9%) 0.2Mb | 372 (5.0%) 0.1Mb |
