## Supplementary material for "Rapid genotyping of tilapia lake virus (TiLV) using Nanopore sequencing": Table S2

**TABLE S2** NanoStat summary statistics of analysis of 274 bp amplicons for each sample (barcode01-05) using the full set of reads and with sub-sampling (sub1K (1000), 500, 100, or 50 reads). ^†^ Coverage or read depth after clustering and filtering steps; ^‡^ mean percent identity of Nanopore raw reads to each sample specific reference; ^§^ during basecalling; BC, barcode

| **BC01** | **Full** | **sub 1k** | **sub 500** | **sub 100** | **sub 50** |
| --- | --- | --- | --- | --- | --- |
| number of reads (coverage) ^†^ | 2276 | 965 | 487 | 98 | 49 |
| Total bases | 634764 | 269359 | 136320 | 27348 | 13600 |
| Total bases aligned | 517537 | 219582 | 110460 | 22297 | 11170 |
| Fraction of bases aligned | 0.8 | 0.8 | 0.8 | 0.8 | 0.8 |
| Median read length | 275 | 275 | 275 | 274 | 274 |
| Mean read length | 278.9 | 279.1 | 279.9 | 279.1 | 277.6 |
| STDEV read length | 11.6 | 11.9 | 13 | 12.5 | 11.5 |
| Read length N50 | 275 | 275 | 275 | 274 | 274 |
| Mean percent identity ^‡^ | 93.4 | 93.3 | 93.3 | 93.5 | 93.2 |
| Mean percent raw reads error ^§^ | 6.6 | 6.7 | 6.7 | 6.5 | 6.8 |
| Median percent identity | 93.6 | 93.6 | 93.6 | 93.7 | 93.2 |
| Mean read quality | 12.4 | 12.4 | 12.4 | 12.5 | 12.3 |
| Median read quality | 12.2 | 12.2 | 12.2 | 12.4 | 11.9 |
| Top 5 longest reads and their mean basecall (quality score) |  |  |  |  |  |
| 1 | 320 (11.7) | 320 (11.4) | 320 (12.8) | 318 (13.6) | 317 (10.9) |
| 2 | 320 (11.6) | 320 (13.5) | 320 (12.0) | 317 (11.4) | 316 (10.0) |
| 3 | 320 (10.4) | 320 (12.8) | 320 (10.2) | 317 (10.9) | 313 (10.6) |
| 4 | 320 (10.1) | 320 (12.0) | 319 (12.4) | 316 (12.0) | 309 (10.5) |
| 5 | 320 (12.2) | 320 (10.2) | 319 (10.8) | 316 (10.0) | 286 (10.8) |
| Top 5 highest mean basecall quality scores and their read lengths |  |  |  |  |  |
| 1 | 18.7 (277) | 18.7 (277) | 18.0 (271) | 17.7 (272) | 16.3 (272) |
| 2 | 18.2 (274) | 17.7 (271) | 17.7 (271) | 16.3 (272) | 15.9 (274) |
| 3 | 18.0 (271) | 17.7 (272) | 17.7 (272) | 15.9 (279) | 15.6 (273) |
| 4 | 17.9 (275) | 17.6 (273) | 17.0 (278) | 15.9 (274) | 15.2 (278) |
| 5 | 17.8 (277) | 17.2 (270) | 16.8 (271) | 15.6 (273) | 14.8 (275) |
| Number, percentage and megabases of reads above quality cutoffs |  |  |  |  |  |
| >Q5 | 2276 (100.0%) 0.6Mb | 965 (100.0%) 0.3Mb | 487 (100.0%) 0.1Mb | 98 (100.0%) 0.0Mb | 49 (100.0%) 0.0Mb |
| >Q7 | 2276 (100.0%) 0.6Mb | 965 (100.0%) 0.3Mb | 487 (100.0%) 0.1Mb | 98 (100.0%) 0.0Mb | 49 (100.0%) 0.0Mb |
| >Q10 | 2276 (100.0%) 0.6Mb | 965 (100.0%) 0.3Mb | 487 (100.0%) 0.1Mb | 98 (100.0%) 0.0Mb | 49 (100.0%) 0.0Mb |
| >Q12 | 1252 (55.0%) 0.3Mb | 532 (55.1%) 0.1Mb | 270 (55.4%) 0.1Mb | 58 (59.2%) 0.0Mb | 24 (49.0%) 0.0Mb |
| >Q15 | 150 (6.6%) 0.0Mb | 60 (6.2%) 0.0Mb | 33 (6.8%) 0.0Mb | 7 (7.1%) 0.0Mb | 4 (8.2%) 0.0Mb |

| **BC02** | **Full** | **sub 1k** | **sub 500** | **sub 100** | **sub 50** |
| --- | --- | --- | --- | --- | --- |
| number of reads (coverage) ^†^ | 1922 | 949 | 477 | 96 | 49 |
| Total bases | 535970 | 264177 | 132807 | 26844 | 13786 |
| Total bases aligned | 432510 | 213530 | 107197 | 21611 | 11100 |
| Fraction of bases aligned | 0.8 | 0.8 | 0.8 | 0.8 | 0.8 |
| Median read length | 275 | 275 | 275 | 275 | 277 |
| Mean read length | 278.9 | 278.4 | 278.4 | 279.6 | 281.3 |
| STDEV read length | 11.5 | 11.4 | 11 | 12.8 | 13.6 |
| Read length N50 | 275 | 275 | 275 | 275 | 277 |
| Mean percent identity ^‡^ | 93.5 | 93.4 | 93.4 | 93.9 | 94.6 |
| Mean percent raw reads error ^§^ | 6.5 | 6.6 | 6.6 | 6.1 | 5.4 |
| Median percent identity | 93.8 | 93.9 | 93.8 | 94.3 | 94.9 |
| Mean read quality | 12.4 | 12.3 | 12.3 | 12.4 | 12.5 |
| Median read quality | 12.2 | 12.2 | 12.2 | 12 | 12.4 |
| Top 5 longest reads and their mean basecall (quality score) |  |  |  |  |  |
| 1 | 320 (10.4) | 320 (12.0) | 320 (11.6) | 320 (12.0) | 318 (12.9) |
| 2 | 320 (10.1) | 320 (11.6) | 320 (10.4) | 318 (12.9) | 317 (11.4) |
| 3 | 320 (11.1) | 320 (11.0) | 320 (10.7) | 317 (11.4) | 313 (10.5) |
| 4 | 320 (11.3) | 320 (11.3) | 319 (12.4) | 313 (10.4) | 312 (12.5) |
| 5 | 320 (11.0) | 320 (10.4) | 319 (12.2) | 313 (10.5) | 310 (10.7) |
| Top 5 highest mean basecall quality scores and their read lengths |  |  |  |  |  |
| 1 | 18.7 (277) | 18.1 (271) | 18.1 (271) | 16.7 (275) | 15.9 (275) |
| 2 | 18.1 (271) | 17.1 (274) | 17.1 (274) | 15.9 (275) | 15.1 (271) |
| 3 | 18.0 (275) | 17.0 (275) | 16.7 (275) | 15.6 (270) | 15.0 (274) |
| 4 | 17.8 (274) | 16.8 (272) | 16.5 (275) | 15.5 (273) | 14.7 (274) |
| 5 | 17.5 (277) | 16.7 (275) | 16.5 (276) | 15.1 (271) | 14.6 (277) |
| Number, percentage and megabases of reads above quality cutoffs |  |  |  |  |  |
| >Q5 | 1922 (100.0%) 0.5Mb | 949 (100.0%) 0.3Mb | 477 (100.0%) 0.1Mb | 96 (100.0%) 0.0Mb | 49 (100.0%) 0.0Mb |
| >Q7 | 1922 (100.0%) 0.5Mb | 949 (100.0%) 0.3Mb | 477 (100.0%) 0.1Mb | 96 (100.0%) 0.0Mb | 49 (100.0%) 0.0Mb |
| >Q10 | 1922 (100.0%) 0.5Mb | 949 (100.0%) 0.3Mb | 477 (100.0%) 0.1Mb | 96 (100.0%) 0.0Mb | 49 (100.0%) 0.0Mb |
| >Q12 | 1068 (55.6%) 0.3Mb | 511 (53.8%) 0.1Mb | 261 (54.7%) 0.1Mb | 47 (49.0%) 0.0Mb | 26 (53.1%) 0.0Mb |
| >Q15 | 100 (5.2%) 0.0Mb | 47 (5.0%) 0.0Mb | 25 (5.2%) 0.0Mb | 6 (6.2%) 0.0Mb | 3 (6.1%) 0.0Mb |

| **BC03** | **Full** | **sub 1k** | **sub 500** | **sub 100** | **sub 50** |
| --- | --- | --- | --- | --- | --- |
| number of reads (coverage) ^†^ | 3485 | 986 | 493 | 99 | 49 |
| Total bases | 970235 | 274952 | 137576 | 27666 | 13771 |
| Total bases aligned | 793561 | 224178 | 112014 | 22479 | 11130 |
| Fraction of bases aligned | 0.8 | 0.8 | 0.8 | 0.8 | 0.8 |
| Median read length | 275 | 275 | 275 | 275 | 275 |
| Mean read length | 278.4 | 278.9 | 279.1 | 279.5 | 281 |
| STDEV read length | 11.1 | 11.5 | 11.4 | 12 | 14 |
| Read length N50 | 275 | 275 | 275 | 275 | 275 |
| Mean percent identity ^‡^ | 93.5 | 93.6 | 93.5 | 93.1 | 93.7 |
| Mean percent raw reads error ^§^ | 6.5 | 6.4 | 6.5 | 6.9 | 6.3 |
| Median percent identity | 93.7 | 94 | 93.7 | 93.3 | 94 |
| Mean read quality | 12.5 | 12.6 | 12.5 | 12.2 | 12.6 |
| Median read quality | 12.3 | 12.4 | 12.4 | 12.2 | 12.5 |
| Top 5 longest reads and their mean basecall (quality score) |  |  |  |  |  |
| 1 | 320 (13.0) | 320 (12.8) | 320 (13.3) | 319 (10.9) | 319 (13.0) |
| 2 | 320 (11.2) | 320 (11.3) | 319 (13.0) | 319 (13.0) | 319 (10.9) |
| 3 | 320 (11.5) | 320 (11.1) | 319 (10.9) | 317 (12.7) | 313 (10.1) |
| 4 | 320 (13.2) | 320 (13.3) | 318 (13.3) | 313 (10.1) | 312 (10.6) |
| 5 | 320 (11.1) | 320 (10.8) | 317 (10.9) | 312 (10.6) | 311 (10.7) |
| Top 5 highest mean basecall quality scores and their read lengths |  |  |  |  |  |
| 1 | 18.8 (272) | 17.6 (273) | 17.5 (277) | 17.2 (280) | 17.2 (280) |
| 2 | 18.1 (274) | 17.6 (272) | 17.2 (280) | 16.7 (272) | 15.7 (271) |
| 3 | 18.1 (271) | 17.4 (273) | 16.9 (270) | 15.7 (271) | 15.6 (275) |
| 4 | 18.1 (271) | 17.4 (275) | 16.8 (274) | 15.6 (275) | 15.2 (273) |
| 5 | 18.0 (275) | 17.2 (280) | 16.7 (272) | 15.2 (273) | 14.8 (275) |
| Number, percentage and megabases of reads above quality cutoffs |  |  |  |  |  |
| >Q5 | 3485 (100.0%) 1.0Mb | 986 (100.0%) 0.3Mb | 493 (100.0%) 0.1Mb | 99 (100.0%) 0.0Mb | 49 (100.0%) 0.0Mb |
| >Q7 | 3485 (100.0%) 1.0Mb | 986 (100.0%) 0.3Mb | 493 (100.0%) 0.1Mb | 99 (100.0%) 0.0Mb | 49 (100.0%) 0.0Mb |
| >Q10 | 3485 (100.0%) 1.0Mb | 986 (100.0%) 0.3Mb | 493 (100.0%) 0.1Mb | 99 (100.0%) 0.0Mb | 49 (100.0%) 0.0Mb |
| >Q12 | 2009 (57.6%) 0.6Mb | 584 (59.2%) 0.2Mb | 287 (58.2%) 0.1Mb | 54 (54.5%) 0.0Mb | 31 (63.3%) 0.0Mb |
| >Q15 | 259 (7.4%) 0.1Mb | 75 (7.6%) 0.0Mb | 35 (7.1%) 0.0Mb | 5 (5.1%) 0.0Mb | 4 (8.2%) 0.0Mb |

| **BC04** | **Full** | **sub 1k** | **sub 500** | **sub 100** | **sub 50** |
| --- | --- | --- | --- | --- | --- |
| number of reads (coverage) ^†^ | 4416 | 998 | 500 | 100 | 50 |
| Total bases | 1231138 | 278535 | 139593 | 27718 | 13894 |
| Total bases aligned | 997426 | 225614 | 112958 | 22605 | 11314 |
| Fraction of bases aligned | 0.8 | 0.8 | 0.8 | 0.8 | 0.8 |
| Median read length | 275 | 275 | 275 | 274 | 274 |
| Mean read length | 278.8 | 279.1 | 279.2 | 277.2 | 277.9 |
| STDEV read length | 12.2 | 11.8 | 11.8 | 10 | 11.7 |
| Read length N50 | 275 | 275 | 275 | 274 | 274 |
| Mean percent identity ^‡^ | 93.5 | 93.6 | 93.4 | 94 | 94 |
| Mean percent raw reads error ^§^ | 6.5 | 6.4 | 6.6 | 6 | 6 |
| Median percent identity | 93.9 | 94 | 93.8 | 94.4 | 94.7 |
| Mean read quality | 12.6 | 12.7 | 12.6 | 12.9 | 12.8 |
| Median read quality | 12.4 | 12.5 | 12.4 | 12.6 | 12.6 |
| Top 5 longest reads and their mean basecall (quality score) |  |  |  |  |  |
| 1 | 320 (13.1) | 320 (12.1) | 320 (13.5) | 319 (11.6) | 319 (11.6) |
| 2 | 320 (11.4) | 320 (11.5) | 320 (12.1) | 317 (11.1) | 317 (11.1) |
| 3 | 320 (11.9) | 320 (13.5) | 319 (11.2) | 315 (11.4) | 315 (11.4) |
| 4 | 320 (12.0) | 319 (11.2) | 319 (14.2) | 309 (13.7) | 303 (10.9) |
| 5 | 320 (12.2) | 319 (13.4) | 319 (11.6) | 303 (10.9) | 293 (10.5) |
| Top 5 highest mean basecall quality scores and their read lengths |  |  |  |  |  |
| 1 | 20.2 (273) | 19.9 (273) | 19.9 (273) | 19.9 (273) | 17.1 (271) |
| 2 | 19.9 (273) | 18.2 (277) | 18.2 (277) | 18.2 (277) | 16.3 (276) |
| 3 | 19.5 (272) | 18.2 (279) | 17.7 (275) | 17.1 (271) | 15.7 (271) |
| 4 | 19.0 (275) | 17.7 (275) | 17.7 (273) | 16.3 (276) | 15.2 (275) |
| 5 | 18.9 (273) | 17.7 (273) | 17.4 (274) | 15.7 (271) | 15.1 (271) |
| Number, percentage and megabases of reads above quality cutoffs |  |  |  |  |  |
| >Q5 | 4416 (100.0%) 1.2Mb | 998 (100.0%) 0.3Mb | 500 (100.0%) 0.1Mb | 100 (100.0%) 0.0Mb | 50 (100.0%) 0.0Mb |
| >Q7 | 4416 (100.0%) 1.2Mb | 998 (100.0%) 0.3Mb | 500 (100.0%) 0.1Mb | 100 (100.0%) 0.0Mb | 50 (100.0%) 0.0Mb |
| >Q10 | 4416 (100.0%) 1.2Mb | 998 (100.0%) 0.3Mb | 500 (100.0%) 0.1Mb | 100 (100.0%) 0.0Mb | 50 (100.0%) 0.0Mb |
| >Q12 | 2612 (59.1%) 0.7Mb | 616 (61.7%) 0.2Mb | 292 (58.4%) 0.1Mb | 66 (66.0%) 0.0Mb | 32 (64.0%) 0.0Mb |
| >Q15 | 379 (8.6%) 0.1Mb | 91 (9.1%) 0.0Mb | 42 (8.4%) 0.0Mb | 10 (10.0%) 0.0Mb | 5 (10.0%) 0.0Mb |

| **BC05** | **Full** | **sub 1k** | **sub 500** | **sub 100** | **sub 50** |
| --- | --- | --- | --- | --- | --- |
| number of reads (coverage) ^†^ | 1354 | 987 | 493 | 100 | 50 |
| Total bases | 380287 | 277033 | 138568 | 28006 | 14090 |
| Total bases aligned | 307780 | 224326 | 111744 | 22619 | 11240 |
| Fraction of bases aligned | 0.8 | 0.8 | 0.8 | 0.8 | 0.8 |
| Median read length | 275 | 275 | 275 | 275 | 275.5 |
| Mean read length | 280.9 | 280.7 | 281.1 | 280.1 | 281.8 |
| STDEV read length | 14.8 | 15 | 16.4 | 12.3 | 13.6 |
| Read length N50 | 275 | 276 | 276 | 275 | 276 |
| Mean percent identity ^‡^ | 93.4 | 93.4 | 93.3 | 93.7 | 93.6 |
| Mean percent raw reads error ^§^ | 6.6 | 6.6 | 6.7 | 6.3 | 6.4 |
| Median percent identity | 93.7 | 93.6 | 93.6 | 94.1 | 94.4 |
| Mean read quality | 12.4 | 12.4 | 12.3 | 12.5 | 12.3 |
| Median read quality | 12.2 | 12.1 | 12.1 | 12.3 | 11.8 |
| Top 5 longest reads and their mean basecall (quality score) |  |  |  |  |  |
| 1 | 320 (11.4) | 320 (12.0) | 320 (13.3) | 318 (11.3) | 318 (11.3) |
| 2 | 320 (11.5) | 320 (11.5) | 320 (11.4) | 315 (11.6) | 315 (11.6) |
| 3 | 320 (13.3) | 320 (10.1) | 320 (12.1) | 313 (10.1) | 312 (10.8) |
| 4 | 320 (12.0) | 320 (10.0) | 320 (11.5) | 312 (10.8) | 312 (10.0) |
| 5 | 320 (11.7) | 320 (12.1) | 319 (10.9) | 312 (10.0) | 309 (10.5) |
| Top 5 highest mean basecall quality scores and their read lengths |  |  |  |  |  |
| 1 | 18.8 (272) | 18.5 (272) | 18.5 (272) | 16.7 (276) | 16.7 (276) |
| 2 | 18.5 (272) | 17.9 (273) | 16.9 (272) | 16.6 (274) | 16.6 (274) |
| 3 | 18.3 (277) | 17.1 (276) | 16.8 (271) | 16.5 (273) | 16.5 (273) |
| 4 | 18.3 (273) | 16.9 (278) | 16.7 (276) | 16.4 (276) | 15.2 (276) |
| 5 | 17.9 (273) | 16.9 (272) | 16.6 (275) | 15.5 (271) | 14.8 (273) |
| Number, percentage and megabases of reads above quality cutoffs |  |  |  |  |  |
| >Q5 | 1354 (100.0%) 0.4Mb | 987 (100.0%) 0.3Mb | 493 (100.0%) 0.1Mb | 100 (100.0%) 0.0Mb | 50 (100.0%) 0.0Mb |
| >Q7 | 1354 (100.0%) 0.4Mb | 987 (100.0%) 0.3Mb | 493 (100.0%) 0.1Mb | 100 (100.0%) 0.0Mb | 50 (100.0%) 0.0Mb |
| >Q10 | 1354 (100.0%) 0.4Mb | 987 (100.0%) 0.3Mb | 493 (100.0%) 0.1Mb | 100 (100.0%) 0.0Mb | 50 (100.0%) 0.0Mb |
| >Q12 | 732 (54.1%) 0.2Mb | 531 (53.8%) 0.1Mb | 265 (53.8%) 0.1Mb | 55 (55.0%) 0.0Mb | 24 (48.0%) 0.0Mb |
| >Q15 | 106 (7.8%) 0.0Mb | 72 (7.3%) 0.0Mb | 33 (6.7%) 0.0Mb | 9 (9.0%) 0.0Mb | 4 (8.0%) 0.0Mb |
