## Supplementary material for "Rapid genotyping of tilapia lake virus (TiLV) using Nanopore sequencing": Table 1

**TABLE 1** Details of TiLV samples used in this study (No. 1-5) whose genomic partial segment 1 sequences were compared with NCBI references (no. 6-19) for phylogenetic analysis

| No. | Sample code | Date | Origin | Fish host | NCBI  Accession no. | References |
| --- | --- | --- | --- | --- | --- | --- |
| 1 | BC01  E-11 cell line day 4 | 2019 | Thailand | Nile tilapia | Not done | This study |
| 2 | BC02  Ti Bang 176-1 | 2017 | Bangladesh | Nile tilapia | Not done | This study |
| 3 | BC03  S1-18 | 2018 | Thailand | RT fingerling | TH-2018-N (MN687745.1) | This study |
| 4 | BC04  m Peru 2018 F3-4 | Feb  2018 | Peru | Nile tilapia | PE-2018_F3-4 (MK425010.1) | This study |
| 5 | BC05  O Peru 2018 F4-5 | Feb  2018 | Peru | Nile tilapia | Not done | This study |
| 6 | IL-2011-Til-4-2011 | May  2011 | Israel | Tilapia | KU751814.1 | (Eyngor et al., 2014)  (Bacharach et al., 2016) |
| 7 | IL-2012-AD-2016 | Aug  2012 | Israel | HT | KU552131.1 | NCBI |
| 8 | TH-2016-TV7 | May  2016 | Thailand | Nile tilapia | KX631936.1 | (Surachetpong et al., 2017) |
| 9 | EC-2012 | Jul  2012 | Ecuador | Nile tilapia | MK392372.1 | (Subramaniam et al., 2019) |
| 10 | TH-2018-K | Aug  2018 | Thailand | NT juvenile | MN687755.1 | (Thawornwattana et al., 2021) |
| 11 | TH-2018-N | Jul  2018 | Thailand | RT fingerling | MN687745.1 | (Thawornwattana et al., 2021) |
| 12 | TH-2019 | Feb  2019 | Thailand | NT fingerlings | MN687765.1 | (Thawornwattana et al., 2021) |
| 13 | PE-2018-F3-4 | Feb  2018 | Peru | Nile tilapia | MK425010.1 | (Pulido et al., 2019) |
| 14 | BD 2017 | Jul  2017 | Bangladesh | Nile tilapia | MN939372.1 | (Chaput et al., 2020) |
| 15 | BD-2017-181 | 2017 | Bangladesh | Nile tilapia | MT466437.1 | (Debnath et al., 2020) |
| 16 | BD-2019E1 | 2019 | Bangladesh | Nile tilapia | MT466447.1 | (Debnath et al., 2020 |
| 17 | BD-2019-E3 | 2019 | Bangladesh | Nile tilapia | MT466457.1 | (Debnath et al., 2020 |
| 18 | USA-2019-WVL19054 | 2019 | USA | Nile tilapia | MN193523-1 | (Al-Hussinee et al., 2018) |
| 19 | USA-2019-WVL19031 | Nov  2018 | USA | Nile tilapia | MN193513.1 | (Al-Hussinee et al., 2018) |

Abbreviations: BC, barcode from Nanopore barcoding kit; country codes: TH, Thailand; IL, Israel; EC, Ecuador; PE, Peru; and BD, Bangladesh. Animal codes: RT, red tilapia (*Oreochromis* spp.); NT, Nile tilapia (*Oreochromis niloticus*); HT, hybrid tilapia (*Oreochromis niloticus x Oreochromis aureus*). Note that samples No.3 and No.4 originated from the same fish specimens used to generate NCBI Sanger references TH-2018-N (No. 11) and PE-2018_F3-4 (No. 13), respectively.
