## Supplementary material for "Rapid genotyping of tilapia lake virus (TiLV) using Nanopore sequencing": Table 2

**TABLE 2** BlastN results of (A) 577 bp consensus sequences generated from the first round PCR products; (B) 231 bp consensus sequences generated from the second semi-nested round PCR products

(A)

| Barcode samples | ^†^Query length (bp) | NCBI top BlastN Hit  TiLV isolate / accession number | % Identity |
| --- | --- | --- | --- |
| BC01 | 577 | TH-2018-N / [MN687745.1](https://www.ncbi.nlm.nih.gov/nucleotide/MN687745.1?report=genbank&log$=nuclalign&blast_rank=1&RID=1AC2SBZ5013) | 100 (577/577 bp) |
| BC02 | 577 | BD-2017-181 / [MT466437.1](https://www.ncbi.nlm.nih.gov/nucleotide/MT466437.1?report=genbank&log$=nucltop&blast_rank=1&RID=1AB7491M013) | 99.83 (575/576 bp) |
| BC03^‡^ | 577 | TH-2018-N / [MN687745.1](https://www.ncbi.nlm.nih.gov/nucleotide/MN687745.1?report=genbank&log$=nucltop&blast_rank=1&RID=1ACAVZAP013) | 100 (577/577 bp) |
| BC04^‡^ | 577 | PE-2018-F3-4 / [MK425010.1](https://www.ncbi.nlm.nih.gov/nucleotide/MK425010.1?report=genbank&log$=nucltop&blast_rank=1&RID=1ACT33NW016) | 100 (577/577 bp) |
| BC05 | 577 | PE-2018-F3-4 / [MK425010.1](https://www.ncbi.nlm.nih.gov/nucleotide/MK425010.1?report=genbank&log$=nucltop&blast_rank=1&RID=1ACT33NW016) | 99.83 (576/577 bp) |

(B)

| Barcode samples | ^†^Query length (bp) | NCBI top BlastN Hit  TiLV isolate / accession number | % Identity |
| --- | --- | --- | --- |
| BC01 | 231 | TH-2018-N / [MN687745.1](https://www.ncbi.nlm.nih.gov/nucleotide/MN687745.1?report=genbank&log$=nuclalign&blast_rank=1&RID=1AC2SBZ5013) | 100 (231/231 bp) |
| BC02 | 231 | BD-2017-181 / [MT466437.1](https://www.ncbi.nlm.nih.gov/nucleotide/MT466437.1?report=genbank&log$=nucltop&blast_rank=1&RID=1AB7491M013) | 100 (230/230 bp) |
| BC03^‡^ | 231 | TH-2018-N / [MN687745.1](https://www.ncbi.nlm.nih.gov/nucleotide/MN687745.1?report=genbank&log$=nucltop&blast_rank=1&RID=1ACAVZAP013) | 100 (231/231 bp) |
| BC04^‡^ | 231 | PE-2018-F3-4 / [MK425010.1](https://www.ncbi.nlm.nih.gov/nucleotide/MK425010.1?report=genbank&log$=nucltop&blast_rank=1&RID=1ACT33NW016) | 100 (230/230 bp) |
| BC05 | 231 | PE-2018-F3-4 / [MK425010.1](https://www.ncbi.nlm.nih.gov/nucleotide/MK425010.1?report=genbank&log$=nucltop&blast_rank=1&RID=1ACT33NW016) | 100 (230/230 bp) |

^†^ Query length of medaka consensus sequences with the primer-binding sites trimmed; ^‡^ Samples previously Sanger sequenced; BC, barcode. For (B) note that for all samples, the BlastN results of consensus sequences (231 bp) generated from sub-sampling (1k, 500, 100, 50 reads) were the same as the ones from no-subsampling.
